# Automated detection and classification of morphologically similar weevil species using AI for precision pest monitoring

**DOI:** 10.64898/2026.11.17.767224

**Authors:** Adama R. Tukuli, Lindsey C. Perkin, Tyler J. Raszick, Xanthe A. Shirley, Zachary P. Cohen, Charles P.-C. Suh

**Affiliations:** USDA-ARS Southern Plains Agricultural Research Center, College Station, Texas, United States of America; USDA-APHIS-PPQ Science & Technology, Insect Management and Molecular Diagnostics Laboratory, Edinburg, Texas, United States of America; USDA-APHIS-PPQ College Station, Texas, United States of America

## Abstract

Accurate identification of morphologically similar insects is important for agricultural surveillance but remains challenging when large and balanced image datasets are limited. We developed Rostrum Recognition: A Weevil Identification Tool (RRWIT), a lightweight object-detection framework for distinguishing 14 weevil taxa, including two economically important weevil species, the boll weevil and pecan weevil, using more than 16,000 expert-verified annotated images. RRWIT was developed by transfer learning from a pretrained YOLO11n detector using a stratified data partition with the test set withheld from model and threshold selection. We evaluated this design with matched random-initialization and transfer-learning experiments, quantitative EigenGradCAM attribution analysis, targeted architectural modifications, and a frozen eight-framework benchmark across three independent training seeds. In the cross-framework benchmark, RRWIT achieved a mean average precision at an intersection-over-union threshold of 0.50 (mAP50) of 0.938 ± 0.005 on held-out test partitions subset, whereas the newer lightweight YOLO26n achieved the highest value, 0.976 ± 0.010, with a similarly compact computational footprint. Post-training 8-bit integer (INT8) quantization reduced the serialized RRWIT model from 10.09 to 2.91 MiB (71.2%) with a small validation mAP50 change from 0.978 to 0.971 but did not accelerate end-to-end inference under the tested central processing unit configuration. RRWIT therefore provides an extensible foundation for automated weevil identification while highlighting the importance of representative biological data, independent external validation, evolving lightweight architectures, and hardware-specific deployment optimization in future development. A publicly accessible RRWIT web application supporting image upload and live image capture for automated weevil detection and classification is available at https://huggingface.co/spaces/adamartukuli/RRWIT.

## 1 Introduction

Curculionidae (Coleoptera), the “true weevils”, is among the most diverse insect families, with more than 6,000 genera and 97,000 described species [1]. Their diversification is closely associated with flowering plants, and many species have a characteristic rostrum bearing modified mouthparts used in feeding and oviposition [1,2]. This diversity includes major agricultural pests, including the pecan weevil, *Curculio caryae* (Horn), which is a serious pest of pecan production and is monitored with Circle traps (Great Lake IPM, Vestaburg, MI) to guide management [3–7], and the boll weevil, *Anthonomus grandis grandis* Boheman, a pest of commercial cotton and is monitored with Pheromone Lure equipped traps (Technical Precision Plastics, Mebane, NC) to guide management, remains subject to surveillance and eradication activities in the Lower Rio Grande Valley of Texas along the U.S.-Mexico border [8,9].

A persistent challenge in pecan and boll weevil surveillance is the frequent capture of morphologically similar, non-pest weevil species in monitoring traps. Closely related taxa can share body size, coloration, overall shape, and other external characters, while reliable species identification may depend on subtle morphological differences requiring specialized taxonomic expertise. Incorrect identification of a non-pest weevil as an economically important pest can trigger unnecessary remedial actions, leading to excess pesticide use, disrupted trade, and increased operational costs. Conversely, incorrect identification of an economic pest as a non-pest can lead to significant losses in yield or increased costs due to treatment delays.

Therefore, accurate, rapid, and field-deployable identification tools are critically needed to improve management decisions and to reduce the costs associated with pest misclassification while reserving uncertain cases for specialist examinations. Moreover, no existing AI models have been specifically tailored to distinguish between pest and non-pest weevils that are morphologically very similar and captured as by-catch in monitoring traps. Deep learning has been applied to cotton pests, boll-weevil classification under limited-data conditions, and other fine-grained insect-identification problems [10–12]. Image classification predicts an image-level class, whereas object detection jointly predicts object identity and spatial extent, usually as a bounding box [13]. For monitoring images that may contain small insects, overlapping targets, debris, or heterogeneous backgrounds, localization and classification are therefore complementary requirements [14,15].

Agricultural detectors have addressed these challenges with feature fusion, attention, and multi-scale processing, including AF-RCNN, AgriPest-YOLO, and Improved Pest-YOLO [16–18]. Deployment-oriented studies further show that predictive accuracy must be considered together with model size, computation, and measured hardware performance [19,20]. Model interpretation adds another diagnostic dimension; attribution maps can indicate where model-associated evidence is concentrated, but they do not establish that a model has learned causal biological characters. Recent lightweight agricultural studies increasingly combine comparative benchmarking, ablation, computational characterization, and attribution visualization rather than relying on a single accuracy value [21].

These considerations motivated development of Rostrum Recognition: A Weevil Identification Tool (RRWIT) by fine-tuning a nano-scale You Only Look Once version 11 detector (YOLO11n) [22] pretrained on Common Objects in Context (COCO) [31] for 14 morphologically similar pest and non-pest weevil taxa. Considering the dataset we have, we hypothesized that transfer learning with a nano-scale one-stage detector would provide a useful balance of fine-grained detection performance and computational efficiency for our finite biological dataset. We tested that design by isolating the effect of pretrained initialization, quantifying EigenGradCAM attribution localization, screening targeted architecture variants, benchmarking eight representative detector frameworks under a common validation/test contract, examining species-specific errors, and measuring model complexity, quantization behavior, and controlled central processing unit (CPU) inference. The study was designed to evaluate RRWIT as a reproducible and extensible framework for automated weevil identification, not to treat the current web prototype as a validated final field-deployment system.

## 2 Materials and methods

### 2.1 Image acquisition and dataset composition

Fourteen weevil taxa, including both pests and non-pests, were selected based on their morphological similarity and frequency of capture in boll weevil and pecan weevil monitoring traps. These included *A. grandis grandis* (boll weevil) and eight non-pest weevil species, *A. g. thurberiae* Pierce*, A. eugenii* Cano, *A. texanus* Dietz, *A. hunteri* Burke and Cate, *A. fulvus* LeConte, *A. cognatus* Burke, *A. peninsularis* Dietz, and *Conotrachelus seniculus* LeConte.

Likewise, five key species included the pecan weevil (*C. caryae*) set – the target pest species and four non-pest weevils: *C. fulvus* Chittenden*, C. proboscideus* Fabricius*, C. pardalis* Chittenden, and *Conotrachelus naso* LeConte.

Specimens were obtained with assistance from the Texas Boll Weevil Eradication Foundation, Texas Department of Agriculture, Texas Pecan Growers Association, and cotton and pecan producers across Texas. Additional contributions were made by the Texas A&M University Insect Collection. Images were collected under diverse conditions using tablets (Samsung Galaxy S9^+^) and a camera mounted to a microscope (Olympus SZ40 with AmScope camera). These devices were selected to simulate both field and laboratory environments, thereby enhancing the generalizability of the dataset. Each specimen was photographed under varying background and lighting conditions. Additional images were obtained from Weevils of Texas, Weevils of North America, iNaturalist, and BugGuide and were taxonomically verified before inclusion [23–26]. Image and specimen metadata were managed with Field Book [27] to ensure meticulous tracking and maintain data integrity throughout the project by assigning each weevil image a unique identifier. The curated collection contained more than 16,000 annotated images distributed among the training, validation, and test partitions while keeping images from the same biological specimen within unique subsets to avoid potential data leakage.

### 2.2 Image preprocessing, annotation, and dataset partitioning

Images from heterogeneous source formats were standardized to JPEG with Pillow [28]. Initial object annotations were generated with Grounding DINO, an open-set detector that links visual and language representations [29]. Every generated bounding box was then inspected and manually corrected in Roboflow annotation platform [30]. Corrections included box position and dimensions, deletion of erroneous boxes, addition of missed objects, and verification of taxonomic labels. The resulting expert-reviewed boxes and labels constituted the ground truth.

The annotated images were partitioned once before model development by taxon-stratified sampling into training (80%), validation (10%), and test (10%) subsets (S1 Fig); partition membership was retained for all subsequent analyses. Training images alone were used for parameter estimation and stochastic augmentation. Augmentations were generated at training during optimization rather than as pre-generated images, so augmented copies were never distributed across partition subsets. Validation data were used for training monitoring, early stopping, checkpoint selection, architecture screening, and, in the later framework benchmark, model-specific confidence-threshold selection. The test partition was not used for fitting or selection. Before the final benchmark accessed test annotations, all checkpoints, thresholds, and evaluator logic were frozen from validation data, and the evaluator was required to reproduce validation metrics while technically unable to open the with-held test annotation subset.

### 2.3 Development of RRWIT

RRWIT was developed by fine-tuning YOLO11n [22] from the Ultralytics yolo11n.pt checkpoint pretrained on the COCO object-detection dataset [31]. The original run used random seed 0 with deterministic training enabled. The archived configuration specified up to 1,000 epochs, early-stopping patience of 15 epochs, batch size 16, 640 × 640-pixel input, six data-loader workers, caching, the complete training partition, and automatic mixed-precision training.

Optimizer selection was performed using the Ultralytics automatic optimizer routine intentionally to ensure reproducibility and consistency. The archived training configuration specified an initial learning rate (lr0) of 0.01, a final learning-rate factor (lrf) of 0.01, momentum of 0.937, weight decay of 0.0005, three warm-up epochs, an initial warm-up momentum of 0.8, and an initial warm-up bias learning rate of 0.1. Because the Ultralytics automatic optimizer routine can internally select the optimizer and override some user-specified optimization parameters, these values are reported as the archived configuration settings rather than as the effective optimizer settings used during training. Loss weights were set to 7.5 for box regression, 0.5 for classification, and 1.5 for distribution focal loss. No layers were frozen; dropout was 0; rectangular and multi-scale training were disabled.

#### 2.3.1 Training-time augmentation

Stochastic augmentation was restricted to the training partition. The seed-0 configuration used hue = 0.015, saturation = 0.7, brightness/value = 0.4, translation = 0.1, scale = 0.5, horizontal-flip probability = 0.5, and Mosaic probability = 1.0, with Mosaic disabled during the final 10 epochs. Rotation, shear, perspective transformation, vertical flipping, MixUp, copy-paste, BGR channel swapping, and multi-scale training were disabled. Validation and test images were evaluated without stochastic training augmentation.

Training used an NVIDIA A100-SXM4-80GB graphics processing unit (GPU) on the USDA Agricultural Research Service SCINet high-performance-computing resource. The selected checkpoint was frozen as RRWIT.v1.0.0.pt (referred as RRWIT in this manuscript) and served as the seed-0 model for diagnostic analyses and prototype deployment. A demonstration web application interface was implemented with Gradio and hosted on Hugging Face Spaces [32,33].

### 2.4 Random-initialization and variance-matched transfer-learning experiment

To isolate pretrained initialization from detector architecture, YOLO11n models initialized from random weights were compared with models initialized from the same frozen COCO-pretrained yolo11n.pt source. Both conditions used the fixed dataset partitions and prespecified seeds 42, 123, and 2025. Random-initialization models were instantiated directly from yolo11n.yaml without loading pretrained weights. Transfer-learning replicates were initialized independently from yolo11n.pt rather than from the seed-0 RRWIT checkpoint. The principal RRWIT training contract was retained, including the 1,000-epoch maximum, 640 × 640 input, batch size 16, patience 15, deterministic execution, automatic mixed precision and optimizer selection, weight decay, warm-up, loss weights, Mosaic schedule, horizontal flipping, and disabled MixUp. Each run was an independent training replicate; validation selected the checkpoint, and test evaluation occurred only afterward. Results were summarized as arithmetic mean ± sample standard deviation (SD) across seeds.

### 2.5 Quantitative EigenGradCAM analysis

The frozen RRWIT checkpoint was examined using EigenGradCAM implemented in pytorch-grad-cam [34]. The purpose of this explainable artificial intelligence (XAI) analysis was to quantify attribution localization relative to the annotated insect and determine whether attribution localization varied with object size or prediction outcome. Inference used an input resolution of 640 × 640 pixels, a confidence threshold of 0.25, and a non-maximum-suppression intersection over union (IoU) threshold of 0.70. For this audit, a prediction was considered correct only when the predicted species was correct and the detection matched the relevant ground-truth bounding box at IoU ≥0.50. Class-targeted EigenGradCAM attribution was calculated from YOLO11n model layer 16, corresponding to the high-resolution P3-scale feature pathway [34–37].

The test partition contained 1,693 images, of which 1,685 contained a single annotated object and formed the audit population. Each image was classified as a correct prediction, species misclassification, localization failure, or no detection under the frozen evaluation rules. Ground-truth object size was calculated as bounding-box area divided by image area and categorized as tiny (<1%), small (1–<10%), medium (10–<50%), or large (≥50%). The complete 1,685-image audit was used to characterize prediction outcomes independently of the attribution subset. Because EigenGradCAM required an available detection, no-detection cases remained in the performance audit but were excluded from attribution calculations.

To preserve error diversity while limiting dominance by the more abundant correct predictions, all qualifying detected incorrect cases were retained, whereas correctly detected cases were ranked by prediction confidence within each species-by-size stratum and limited to a maximum of three observations per stratum. This deterministic procedure produced a fixed attribution subset of 258 images, comprising 115 correct and 143 incorrect detections. The deliberate enrichment of incorrect predictions was used to compare attribution behavior between successful and unsuccessful predictions and was not used to estimate the test-set error rate. Attribution maps were normalized within each image before quantitative analysis.

Ground-truth-box enrichment was calculated as the fraction of total EigenGradCAM attribution within the annotated bounding box divided by the fraction of image area occupied by that box. An enrichment value of 1 indicates attribution proportional to box area, values >1 indicate relative concentration of attribution within the insect box, and values <1 indicate relative under-enrichment within the box. Because bounding-box area contributes directly to the normalization, enrichment was interpreted jointly with continuous object-area fraction. Representative examples were selected without visual-preference screening: among correctly detected in-house images, the observation closest to the median enrichment within each size stratum was selected, with IoU, prediction confidence, and filename used sequentially as deterministic tie-breakers. Associations between continuous object-area fraction and enrichment were assessed separately for correct and incorrect predictions using Spearman rank correlation, with 5,000 bootstrap replicates used where uncertainty intervals were reported. Attribution maps were interpreted as diagnostic evidence of model attention patterns rather than as evidence of causal taxonomic characters [35–37].

### 2.6 Targeted architecture ablation

EigenGradCAM motivated a controlled test of whether additional shallow-scale information or explicit channel-spatial recalibration improved the RRWIT YOLO11n baseline. For this initial architecture screen, all four models were trained using the same random seed 42 to hold the seed constant while comparing architectural modifications. The four variants were A0, the unmodified COCO-pretrained baseline; A1, a P2-to-P3 feature-assistance pathway; A2, a Convolutional Block Attention Module (CBAM) inserted at P3; and A3, the combined P2-assistance plus P3-CBAM design. The CBAM module sequentially applies channel and spatial attention [38]. All variants retained P3, P4, and P5 detection outputs and were trained using the same fixed training and validation partitions and RRWIT protocol. Selection used mean average precision across intersection-over-union (IoU) thresholds from 0.50 to 0.95 (mAP50-95) as the primary criterion, followed by mean average precision at an IoU threshold of 0.50 (mAP50) and fewer parameters as tie-breakers. The test split was not accessed during screening. Parameter counts were obtained from the exact model implementations, and theoretical computational complexity, expressed as billions of floating-point operations (GFLOPs) at an input resolution of 640 × 640 pixels, was estimated using the Ultralytics get_flops implementation. A modified architecture would advance to multi-seed evaluation and subsequent test-set confirmation only if it surpassed A0 on the prespecified validation criteria.

### 2.7 Eight-framework, three-seed benchmark

RRWIT was compared with seven representative detector configurations: YOLOv5nu, YOLOv8n, YOLO12n, YOLO26n, Single Shot MultiBox Detector Lite (SSDLite320) MobileNetV3-Large, EfficientDet-D0, and Faster R-CNN MobileNetV3-Large-320 feature pyramid network (FPN) [39–46]. Published architecture and implementation characteristics are summarized in S1 Table. The benchmark was designed to compare complete framework-specific detector configurations on the same biological task rather than to isolate architecture under an identical input-resolution and optimization regime. Accordingly, reference or architecture-appropriate input resolutions and family-specific training configurations were retained rather than forcing all detector families to use a single resolution, optimizer, or batch size. The YOLO models used 640 × 640-pixel inputs and batch size 16, SSDLite320 used 320 × 320 pixels and batch size 32, EfficientDet-D0 used 512 × 512 pixels and batch size 16, and Faster R-CNN MobileNetV3-Large-320 FPN used its low-resolution 320-pixel configuration with batch size 8. All frameworks used the same frozen RRWIT source-image partitions and common evaluation definitions. Because input resolution and training configuration can influence both predictive performance and computational requirements, results were interpreted as comparisons among the evaluated framework-specific configurations rather than as an iso-resolution architecture ablation.

Each framework was trained independently with seeds 42, 123, and 2025, producing 24 model-seed combinations on identical source-image partitions and 14 classes. Framework adapters transformed the same annotations into the coordinate and label conventions required by each implementation, and reconstruction audits checked box equivalence. Validation and test preprocessing were deterministic. For each model-seed pair, canonical checkpoint selection used validation COCO-style mAP50-95 as the primary criterion [31]; confidence thresholds required for F1 metrics were also selected using validation predictions. Before test access, a locked gate reloaded every selected checkpoint through its framework-specific inference path and reproduced canonical validation metrics and locked-threshold F1 values within prespecified tolerances. The test-enabled evaluator was authorized only after all 24 gates passed. Test results were not subsequently used to retrain models, substitute checkpoints, adjust thresholds, or select architectures.

### 2.8 Performance metrics and statistical analysis

The primary benchmark metric was COCO-style mean average precision averaged over IoU thresholds 0.50 to 0.95 in increments of 0.05 (mAP50-95) [31]. Additional metrics included mAP at IoU 0.50 (mAP50), mAP at IoU 0.75 (mAP75), and average recall with at most 100 detections per image (AR100). At validation-selected confidence thresholds, precision and recall were used to derive macro, micro, and support-weighted F1 scores; species-specific AP50-95, AP50, and F1 were retained to prevent aggregate metrics from obscuring low-support taxa.

Three-seed experiments were summarized as arithmetic mean ± sample standard deviation (SD; n - 1 degrees of freedom). Because each detector had only three independent training replicates, small between-model differences were interpreted descriptively rather than as strong inferential evidence; no hypothesis test was used to promote a model based on locked-test results. EigenGradCAM area-enrichment associations used Spearman rank correlation and, where reported, 5,000 bootstrap replicates.

### 2.9 Computational footprint analysis

Computational characteristics were measured from the exact trained 14-class models.

Registered model parameters were counted directly, and 32-bit floating-point (FP32) parameter memory was calculated from registered parameter tensors stored in 32-bit floating point, excluding buffers, optimizer/scheduler state, training history, metadata, and serialization overhead. Theoretical computation was estimated for a batch-size-one eager forward pass at each framework’s frozen input resolution with torch.utils.flop_counter.FlopCounterMode [34] and reported as GFLOPs. GFLOPs were treated as an operation-count descriptor, not as latency, throughput, energy use, or hardware-specific speed. Raw framework-native training checkpoint sizes were not compared because their serialized contents were not equivalent across families; deployment artifact size was evaluated separately after export to a common representation.

### 2.10 Deployment-oriented ONNX and INT8 evaluation

Four representative lightweight detectors were carried forward after the accuracy benchmark: YOLO26n seed 123; YOLO12n seed 42; RRWIT seed 2025; and EfficientDet-D0 seed 42. Each checkpoint was selected using validation performance only and was analyzed separately from the three-seed benchmark means. Models were exported in full precision to Open Neural Network Exchange (ONNX) opset 17 [47] and audited for structural validity and output parity. Static post-training 8-bit integer (INT8) quantization used ONNX Runtime [48] with QuantizeLinear/DeQuantizeLinear representation, signed 8-bit activations and weights (S8S8), per-tensor weight quantization, and MinMax calibration. Calibration used a frozen cohort of 512 training images; validation and test images were excluded.

Common-protocol INT8 artifacts were evaluated on validation data for mAP50-95, mAP50, support-weighted F1, proportional mAP50-95 retention, serialized artifact size, and percentage size reduction. A prespecified aggregate accuracy gate required absolute declines ≤0.010 in mAP50-95, mAP50, and weighted F1. Because EfficientDet-D0 degraded substantially under the common S8S8/per-tensor protocol, additional signed/unsigned activation representations and per-channel weight quantization were evaluated as architecture-specific diagnostic sensitivity analyses only; they were not substituted for the authoritative common-protocol result.

#### 2.10.1 Controlled CPU inference

FP32 and common-protocol INT8 artifacts were benchmarked with ONNX Runtime CPUExecutionProvider. The batch-size-one protocol used four intra-operation threads, 16 warm-up images, and three timed repetitions of the same 128 images drawn exclusively from the training partition. Model-core latency measured ONNX graph execution after input preparation; end-to-end latency additionally included image loading/decoding, family-canonical preprocessing, ONNX inference, detector-specific decoding and non-maximum suppression where required, and conversion to the common prediction representation. Median latency, 95th-percentile end-to-end latency, throughput, and peak process memory were retained. Results characterize this x86 CPU/ONNX Runtime configuration and were not extrapolated to ARM processors, neural-processing units, GPUs, or other edge accelerators.

### 2.11 Software environment reproducibility and code availability

The RRWIT model used Python 3.11.7, PyTorch 2.5.1 with CUDA 12.4, and Ultralytics 8.3.85. Architecture screening used Python 3.10.20, PyTorch 2.5.1 with CUDA 12.4, and Ultralytics 8.3.28. The frozen cross-framework benchmark used Python 3.10.20, PyTorch 2.7.1 with CUDA 12.6, Torchvision 0.22.1, Ultralytics 8.4.118, effdet 0.4.1, timm 0.9.16, pycocotools 2.0.11, ONNX 1.22.0, and ONNX Runtime 1.23.2 to accommodate framework-specific dependency constraints. Requirement files and codes used in the main analysis are available at https://github.com/adama960/A-Weevil-Identification-Tool-RRWIT-GitHub repository.

The final fine-tuned YOLOv11n-based model checkpoint is publicly available on our Hugging Face Space.

## 3 Results and Discussion

### 3.1 RRWIT achieves robust overall species-level detection across morphologically similar weevils

Manual annotation review and dataset-level diagnostics showed representation of all 14 taxa together with broad variation in normalized object position and bounding-box dimensions (S1 and S2 Figs). Class support remained uneven: *A. grandis grandis* was the largest class, whereas several taxa had substantially fewer images; most notably *C. fulvus* (S1 Fig).

Bounding-box centers were predominantly specimen-centered but occupied a wider region of the image plane, and width-height distributions covered a broad range. The expected positive width-height association and geometric constraints between box size and valid center locations were consistent with normalized object annotations (S2 Fig). The seed-0 RRWIT training run showed progressive convergence before early stopping at epoch 565 (Fig 1). Training and validation box, classification, and distribution focal losses generally decreased over training (Fig 1A and 1B), while validation mAP50 and mAP50-95 increased and gradually approached a plateau (Fig 1C). The checkpoint at epoch 550 was selected on the basis of validation performance. Although training losses continued to decline, the validation losses did not show pronounced late-stage deterioration, and validation mean average precision remained near its highest levels toward the end of training. Thus, the learning curves did not indicate substantial late-stage overfitting in this run.

**Fig 1.**
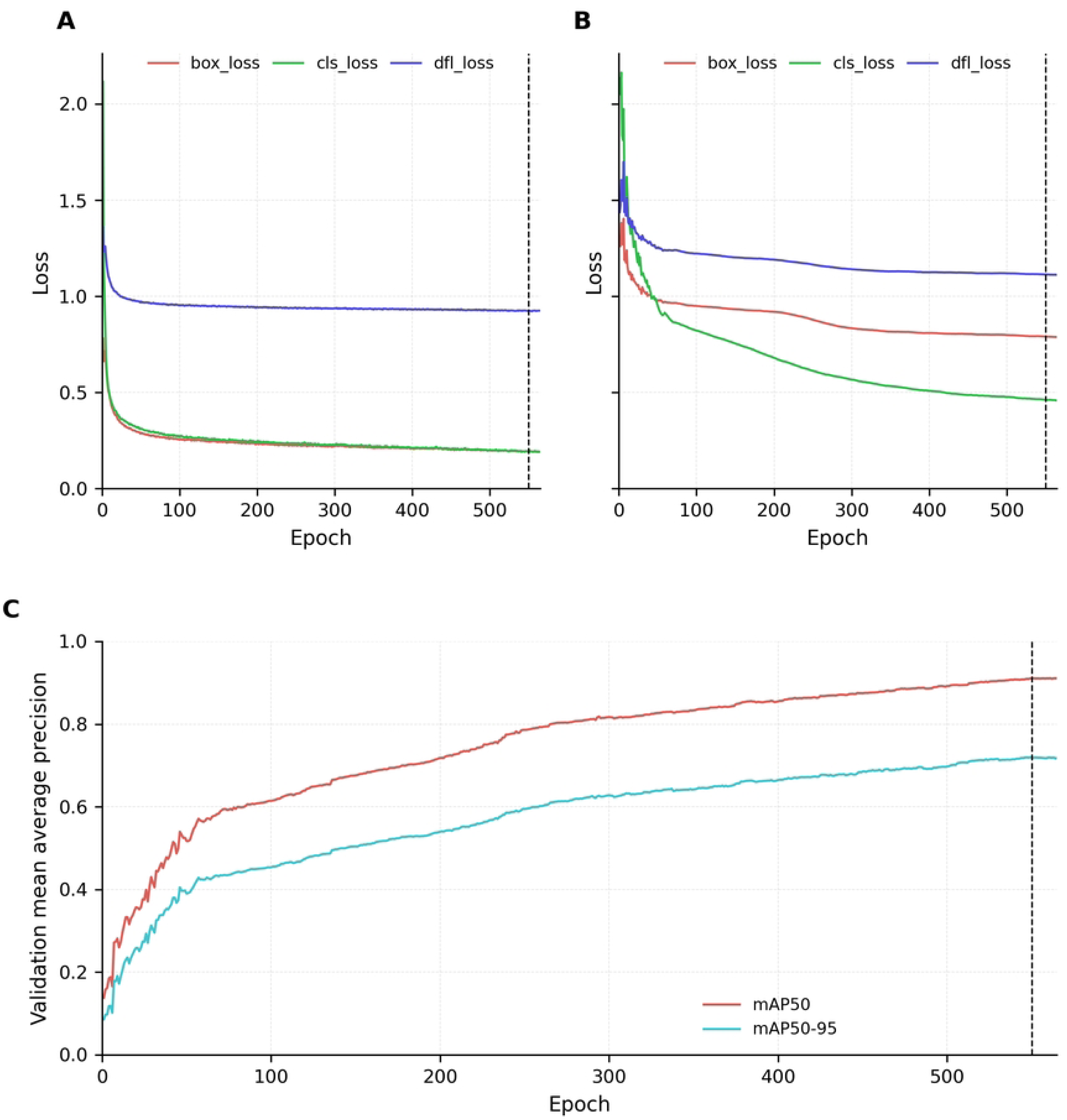
Training convergence and validation performance of Rostrum Recognition: A Weevil Identification Tool (RRWIT). (A) Training bounding-box regression loss (box_loss), classification loss (cls_loss), and distribution focal loss (dfl_loss). (B) Corresponding validation losses. (C) Validation mean average precision at an intersection-over-union (IoU) threshold of 0.50 (mAP50) and mean average precision averaged across IoU thresholds from 0.50 to 0.95 (mAP50-95). The vertical dashed line marks the validation-selected checkpoint at epoch 550; training terminated at epoch 565 under the early-stopping criterion.

Evaluation of the validation and held-out test partitions showed strong overall detection performance, with some reduction in performance on the previously unseen test data (Fig 2). Overall mAP50 was 0.96 on the validation partition and 0.87 on the test partition. The precision-recall curves likewise showed generally strong performance across taxa, although the degree of precision-recall trade-off varied among species (Fig 2C and 2D). The all-class F1 curve reached a maximum of approximately 0.84 at a confidence threshold of 0.464 on validation data and 0.81 at 0.393 on test data (Fig 2E and 2F). Species-level precision, recall, and F1 curves showed that performance was not uniform across taxa, with several classes maintaining comparatively high values across a broad range of confidence thresholds while others showed greater sensitivity to threshold selection (Fig 2A, 2B, and 2E–H). Together, the validation-to-test changes indicate that RRWIT retained substantial discriminatory performance on held-out previously unseen test partitions subset data while also revealing species-specific differences that are obscured by aggregate mAP values.

**Fig 2.**
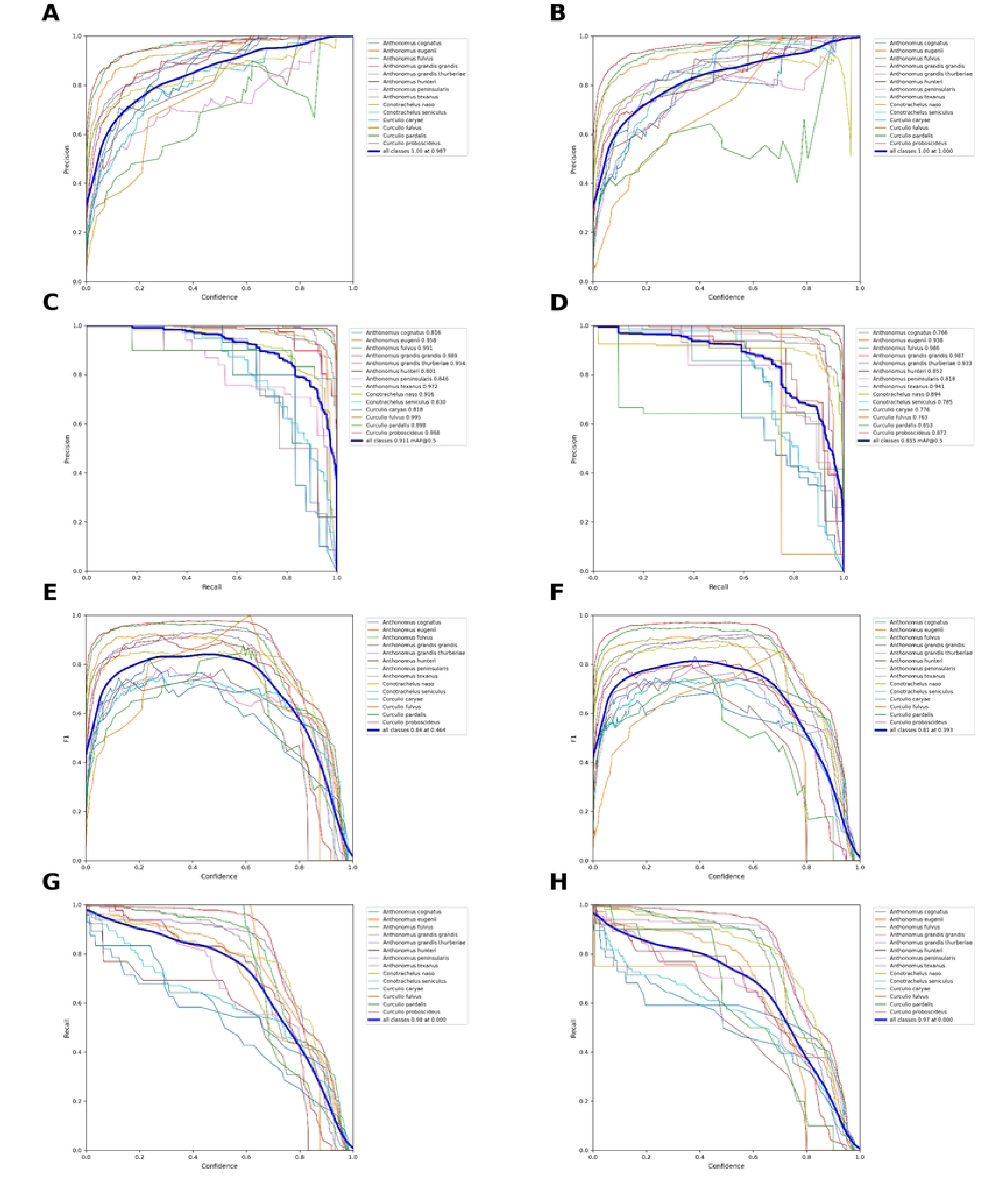
Precision, recall, F1, and precision-recall performance of RRWIT on the validation and held-out test partitions subset data. Left-column panels show validation performance and right-column panels show held-out previously unseen test performance. (A, B) Precision as a function of confidence threshold; (C, D) precision-recall curves; (E, F) F1 score as a function of confidence threshold; and (G, H) recall as a function of confidence threshold. Thin colored lines represent individual weevil taxa, and the thick blue line represents the all-class summary. Values reported with the all-class F1 curves indicate the maximum F1 score and corresponding confidence threshold. Values reported in the precision-recall legends are average precision at an intersection-over-union threshold of 0.50 (AP50), with the thick blue curve reporting mean average precision across classes (mAP50).

The normalized confusion matrices further resolved species-specific differences in RRWIT performance (Fig 3). Correct classifications dominated the diagonal for most taxa in both validation and test data, but the magnitude of the diagonal varied among species. On the validation partition, diagonal proportions ranged from 0.54 to 1.00, with particularly high values for *Curculio fulvus* (1.00), *Anthonomus grandis grandis* (0.98), *Anthonomus texanus* (0.95), and *Anthonomus fulvus* (0.94) (Fig 3A). On the independent held-out previously unseen test partition, diagonal proportions ranged from 0.50 to 0.98, with high values for *Anthonomus grandis thurberiae* (0.98), *Anthonomus grandis grandis* (0.97), *Anthonomus texanus* (0.94), and *Anthonomus fulvus* (0.92) (Fig 3B). Lower diagonal proportions were observed for some taxa, including *Curculio fulvus* (0.50), *Curculio caryae* (0.59), *Anthonomus cognatus* (0.61), *Anthonomus hunteri* (0.62), and *Conotrachelus seniculus* (0.62) on the test partition. The off-diagonal structure shows that errors were concentrated among particular taxon pairs rather than being uniformly distributed across the 14 classes. These class-specific patterns complement the aggregate mAP and F1 results by identifying taxa for which discrimination remained more challenging. This pattern is consistent with the subtle characters used in weevil taxonomy [49,50], although class representation, orientation, illumination, scale, and background can also contribute to errors [16–18].

**Fig 3.**
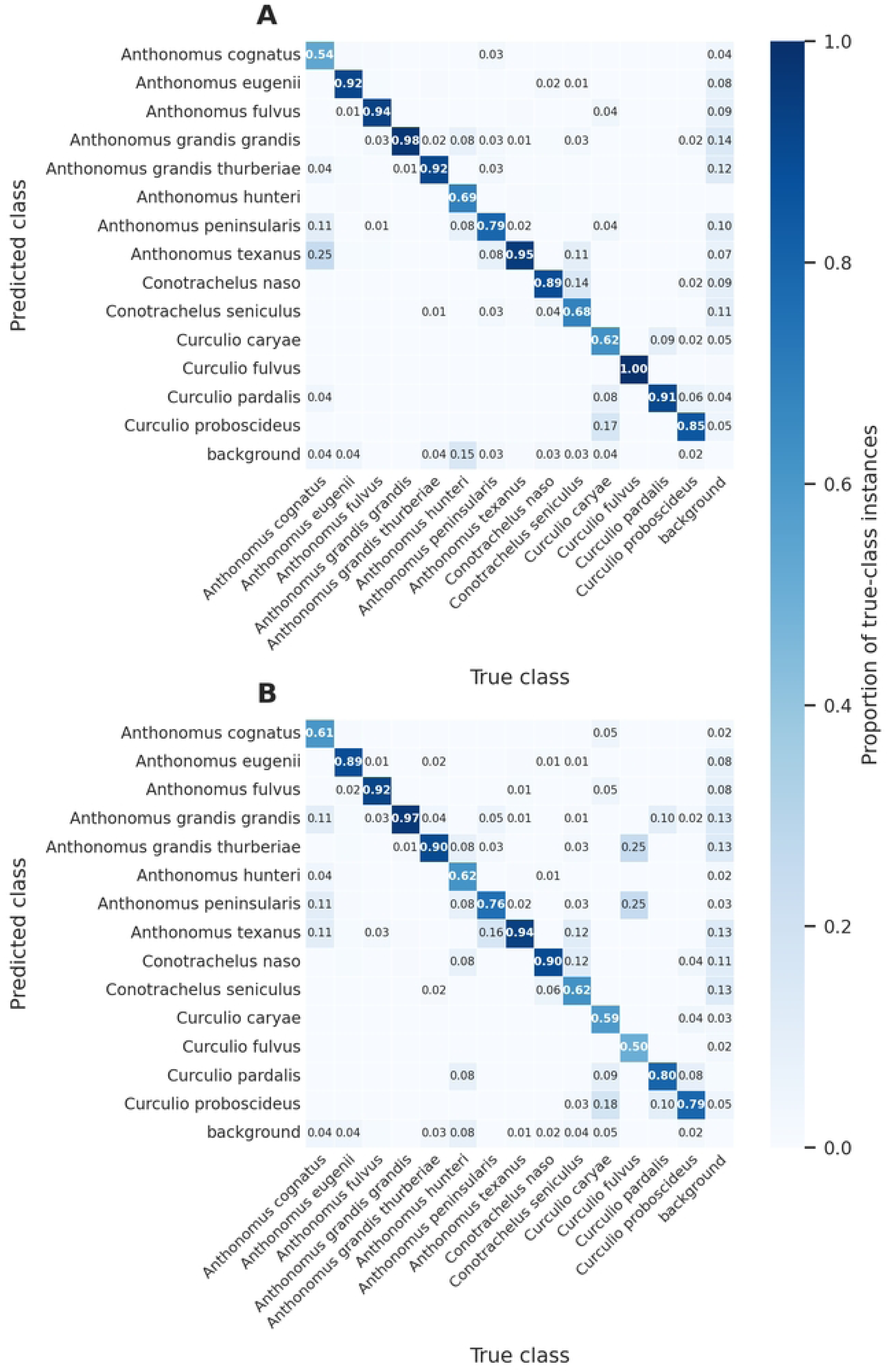
Normalized confusion matrices for RRWIT on the validation and held-out previously unseen test partitions subset data. (A) Validation partition. (B) Independent test partition. Rows represent predicted classes and columns represent true classes. Values are normalized within each true-class column and therefore represent the proportion of instances for each true class assigned to each predicted class under the detection-matching criteria. Diagonal cells indicate correct class assignments, whereas off-diagonal cells indicate misclassifications.

To further evaluate the robustness of the RRWIT model, a qualitative visual analysis was conducted on representative individual and batches of images (Fig 4 and S3 Fig). These results highlight both the localization and classification performance of the model across several morphologically similar species. As shown in Fig 4, ground truth annotations (Fig 4A) reflect the diversity in image acquisition. The corresponding model predictions (Fig 4B) demonstrate that the model reliably delineated bounding boxes and correctly assigned species labels with high confidence across most samples. Colors correspond to species classes, with label confidence shown numerically in the predictions. In the representative batch shown in Fig 4, 15 of 16 images were correctly classified, with one misclassification involving *A. eugenii* (green yellow color), underscoring the model’s capability when species-specific features are distinct and imaging conditions are optimized. When RRWIT is tested with a single image (S3 Fig), it automatically detects the presence of weevils, generates bounding boxes, and assigns a species identification based on the trained detection model. The predicted species label and associated confidence score are displayed directly on the image, allowing the user to visually confirm the result. Overall, the qualitative inspection is consistent with the quantitative evaluations in Figs 1–4 while also illustrating model errors under challenging image conditions. RRWIT demonstrates detection performance under various image conditions, including lighting variation and occlusion. Notable misclassifications are evident in some panels due to species similarity and challenging visual contexts. These results highlight both model strengths and limitations under real-world conditions. It also emphasizes the importance of expanding image diversity and representation, particularly for morphologically ambiguous or underrepresented species.

**Fig 4.**
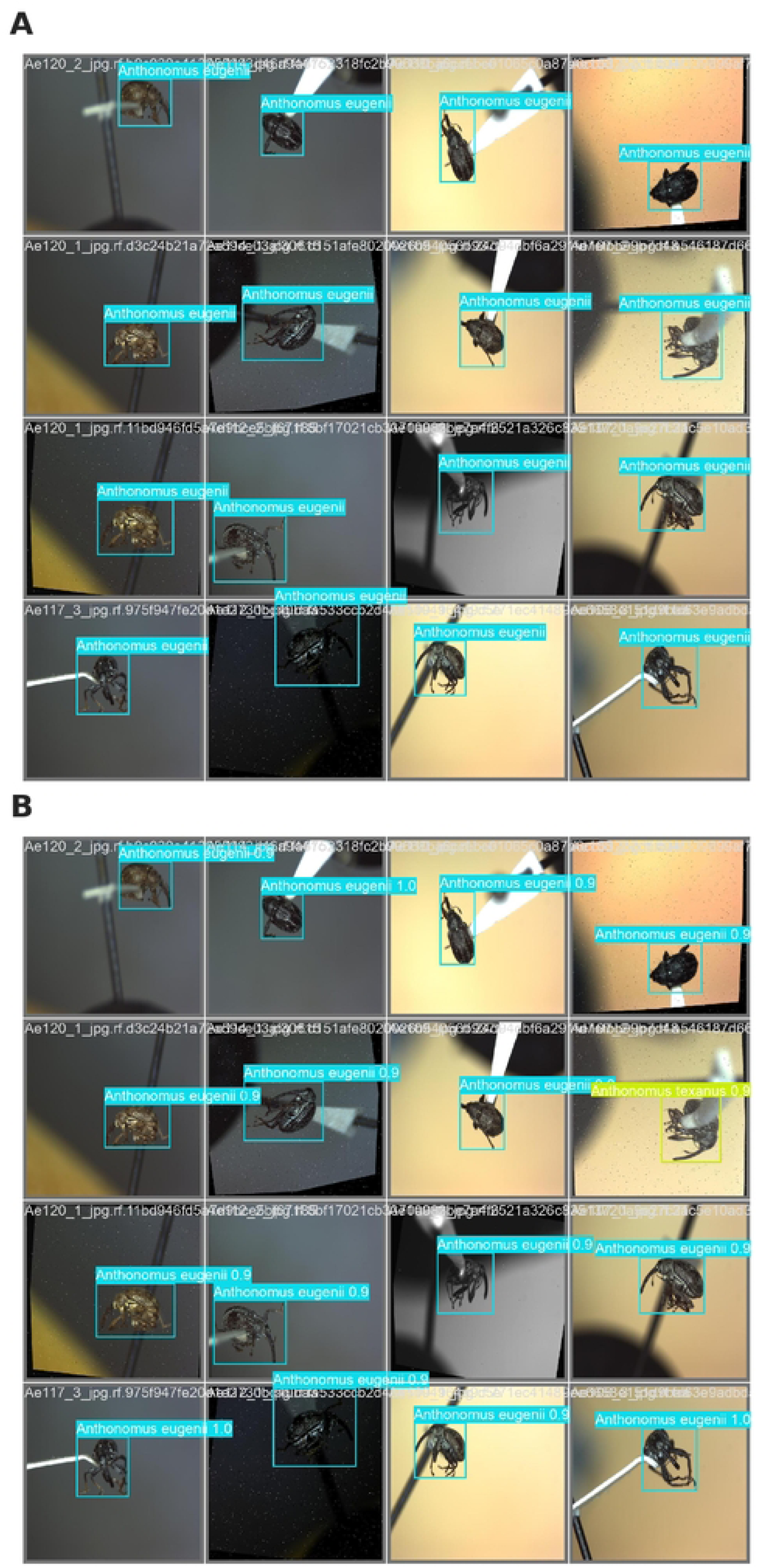
Ground truth vs. RRWIT predictions for one representative batch consisting of 16 images. (A) Displays ground truth labels, while (B) displays predicted bounding boxes and class labels by the RRWIT. Colors correspond to species classes, with label confidence shown numerically in the predictions.

In the broader agricultural context, reported accuracies vary substantially with taxonomic granularity, object scale, annotation policy, and evaluation definition; direct cross-paper numerical ranking can therefore be misleading [51–55]. RRWIT’s main contribution is the application of lightweight object detection to a fine-grained pest/non-pest weevil identification problem, with the subsequent controlled experiments providing the more rigorous basis for comparative conclusions.

Cross-study comparisons provide context but require caution because agricultural detection papers differ in taxonomic granularity, object scale, image acquisition, class balance, dataset partitioning, and evaluation definitions. AF-RCNN reported 56.4% mAP on a 24-class agricultural pest dataset [16], AgriPest-YOLO reported 71.3% mAP on an approximately 25,000-image light-trap dataset [17], and Improved Pest-YOLO reported 73.4% mAP after adding efficient channel attention and transformer-based feature processing [18]. These studies emphasize the difficulty of multi-species pest detection when targets are small, densely distributed, or observed against complex backgrounds; their numerical values should not be treated as direct rankings against RRWIT because the underlying tasks and metrics differ.

Recent work increasingly evaluates accuracy together with computational efficiency. Faster-PestNet was developed as a lightweight crop-pest detector [51], whereas PestLite reported 90.7% mAP50 while reducing a YOLOv5-based model from 7.03 to 6.09 million parameters [52]. A UAV-oriented pest detector evaluated on five broad pest groups reported mean precision of 96.0%, recall of 93.0%, and mAP of 95.0% [53]. Those values exceed the RRWIT test mAP50, but the UAV study separated five relatively broad pest groups, whereas RRWIT distinguishes 14 morphologically similar weevil taxa, including congeneric pest and non-pest lookalikes. Differences in reported accuracy therefore reflect both detector performance and task difficulty.

The distinction between mAP50 and stricter localization metrics is also increasingly explicit in contemporary agricultural vision. A rice-pest study reported 98.8% mAP50 and 78.6% mAP50-95 [54]. YOLO-DCPG, developed for intensive small-target pest detection on Pest24, reported 74.0% mAP50 and 47.5% mAP50-95 with 1.47 million parameters and 5.7 GFLOPs [55]. Together, these studies support reporting multiple complementary metrics rather than presenting a single mAP value as a complete description of detector behavior. RRWIT contributes to this literature from a distinct fine-grained taxonomic perspective, where operational value depends on separating pest species from visually similar by-catch rather than only detecting visually dissimilar pest categories.

A trial RRWIT web application was implemented as a user-facing web interface at https://huggingface.co/spaces/adamartukuli/RRWIT. Users can upload an image or use live image capture and receive predicted bounding boxes, species labels, confidence values, and an adjustable confidence threshold (S1 Video). The interface demonstrates how the trained detector can be presented for interactive expert-assisted identification. It is a functional prototype, not evidence of independent field validation. The present study did not systematically measure application-level latency, user performance, scalability, energy consumption, on-device, performance on representative mobile and edge hardware.

### 3.2 Transfer learning improves detection performance and training consistency

Randomly initialized YOLO11n models learned the task, with declining training and validation losses and increasing validation mAP across all three seeds, although their training trajectories and validation-selected stopping points varied among runs (Fig 5). These results show that random initialization did not prevent optimization of the YOLO11n architecture and provide a matched baseline for evaluating the contribution of pretrained initialization to held-out performance and between-seed variability.

**Fig 5.**
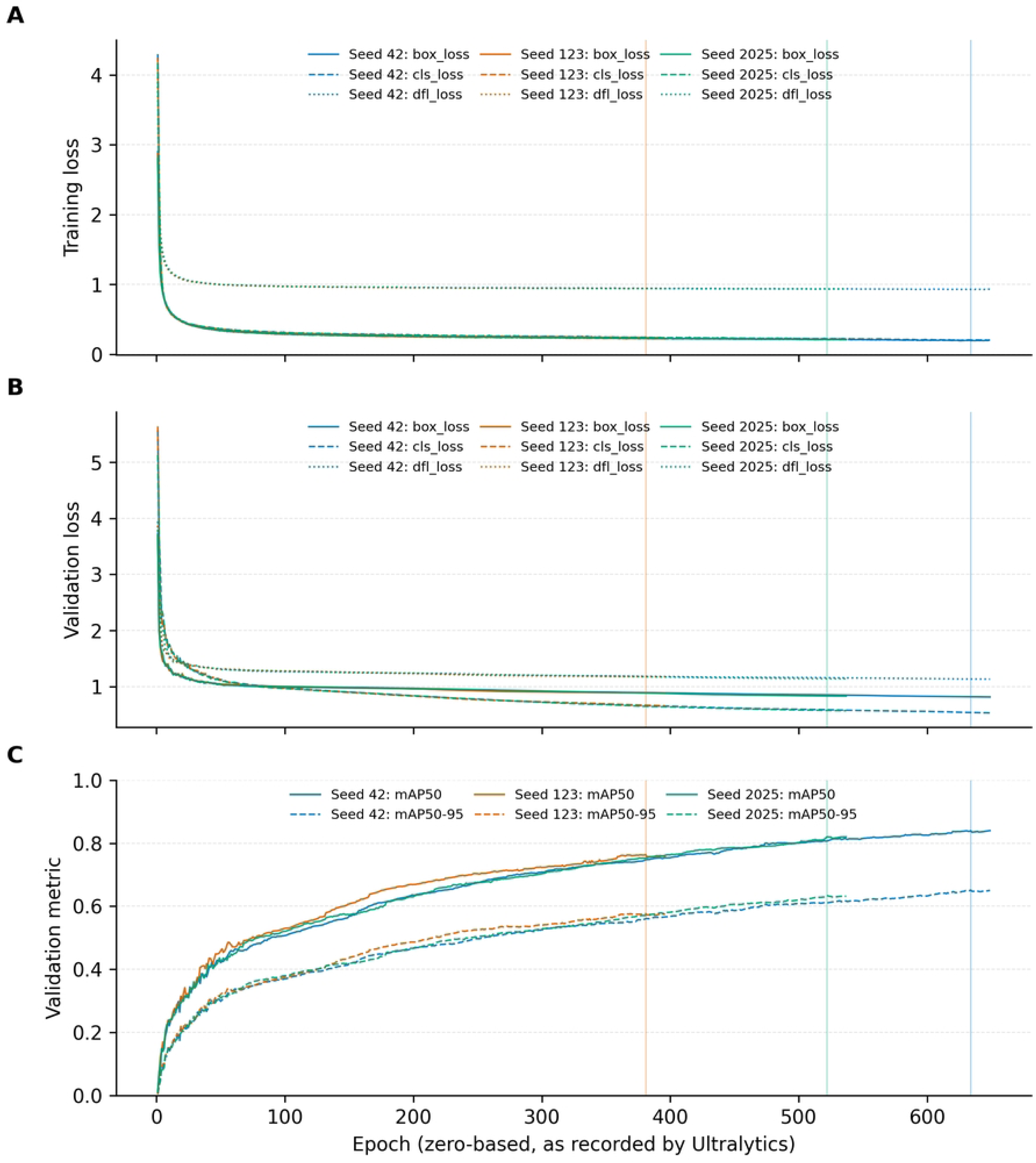
Three-seed training convergence of randomly initialized YOLO11n controls. Seeds 42, 123, and 2025 were trained under the matched RRWIT training protocol. (A) Training box, classification, and distribution focal losses. (B) Corresponding validation losses. (C) Validation mAP50 and mAP50-95. Vertical lines mark the validation-selected checkpoint for each seed.

On the held-out test partition, the randomly initialized models achieved mAP50 values of 0.798, 0.723, and 0.762 for seeds 42, 123, and 2025, respectively, corresponding to a mean ± sample standard deviation (SD) of 0.761 ± 0.038 (Fig 6A). Their corresponding mAP50-95 values were 0.616, 0.550, and 0.590, with a mean of 0.586 ± 0.033. Precision, recall, and F1 also varied among seeds (Fig 6A), while species-level AP50 and AP50-95 showed substantial heterogeneity in both average performance and between-seed variability (Fig 6B). Together, these results demonstrate that the randomly initialized models could learn the detection task but exhibited lower performance and noticeable variation in held-out performance across the three seeds.

**Fig 6.**
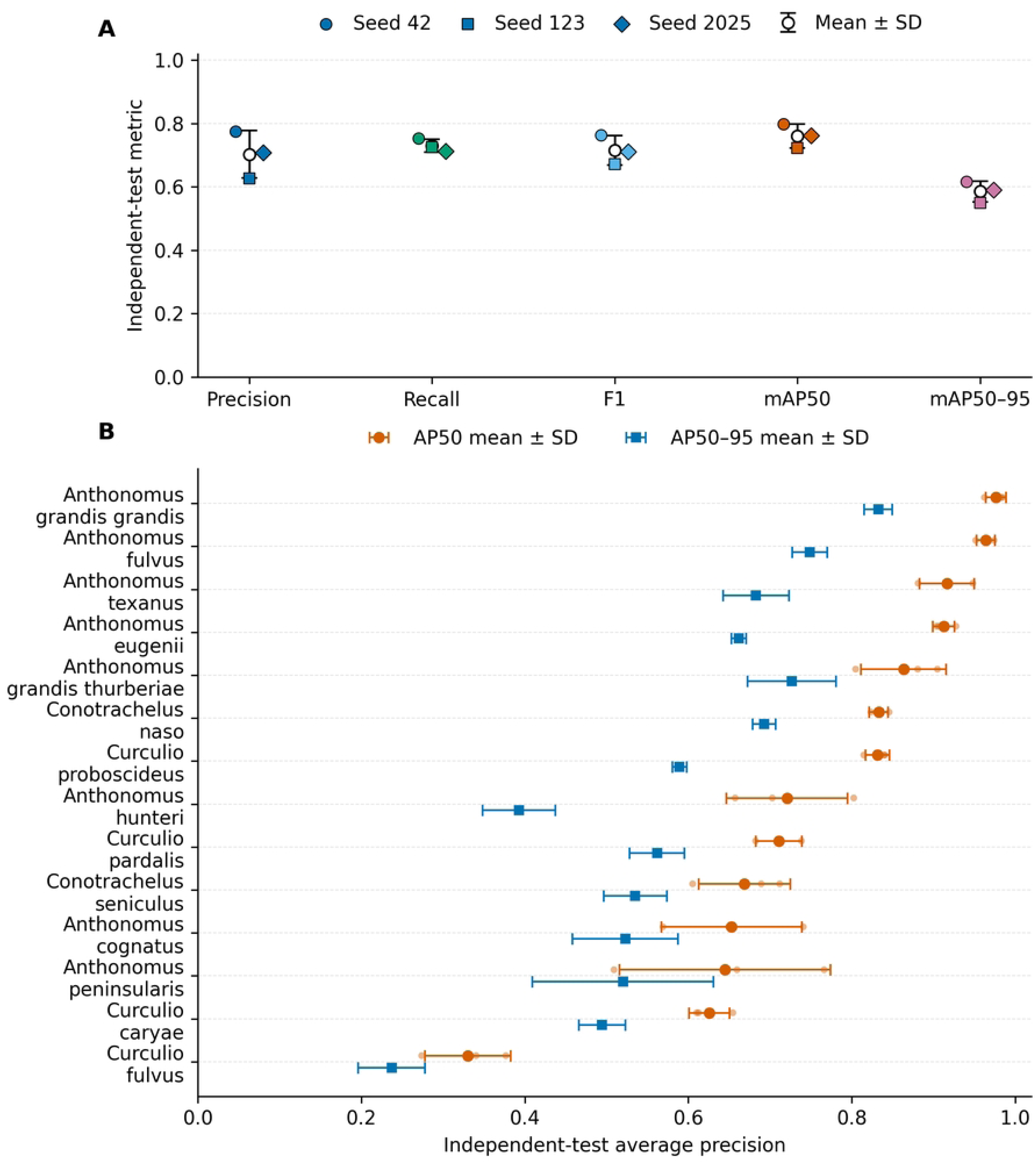
Held-out test performance of three randomly initialized YOLO11n controls. (A) Overall precision, recall, F1, mAP50, and mAP50-95 for seeds 42, 123, and 2025; open symbols and error bars indicate the arithmetic mean ± sample SD across the three seeds. (B) Species-level AP50 and AP50-95 summarized as mean ± sample SD across the same three seeds.

The matched COCO-pretrained YOLO11n models achieved test mAP50 values of 0.803, 0.836, and 0.799 for seeds 42, 123, and 2025, respectively, corresponding to 0.813 ± 0.020, and mAP50-95 values of 0.622, 0.645, and 0.617, corresponding to 0.628 ± 0.015 (Fig 7A). Relative to random initialization, pretrained initialization therefore increased mean mAP50 by 0.052 and mean mAP50-95 by 0.042. Between-seed variability was also lower for the pretrained models; the sample SD decreased from 0.038 to 0.020 for mAP50 and from 0.033 to 0.015 for mAP50-95. The mAP50-95 difference favored pretrained initialization for each matched seed, with the largest difference observed for seed 123, which had the lowest randomly initialized performance. Given the small number of replicated seeds (n = 3 per initialization condition), these differences are interpreted descriptively rather than as evidence of statistical significance. Species-level AP remained heterogeneous after pretrained initialization (Fig 7B), indicating that the overall improvement did not eliminate differences in detection difficulty among taxa. The three replicated pretrained models did not reproduce the performance of the seed-0 RRWIT checkpoint, illustrating the stochastic variability associated with model training. The seed-0 checkpoint was retained as the deployed RRWIT model, whereas the subsequent three-seed experiment was designed as an independent, matched comparison of pretrained and randomly initialized models and therefore was interpreted on the basis of paired initialization conditions rather than comparison with seed-0 RRWIT.

**Fig 7.**
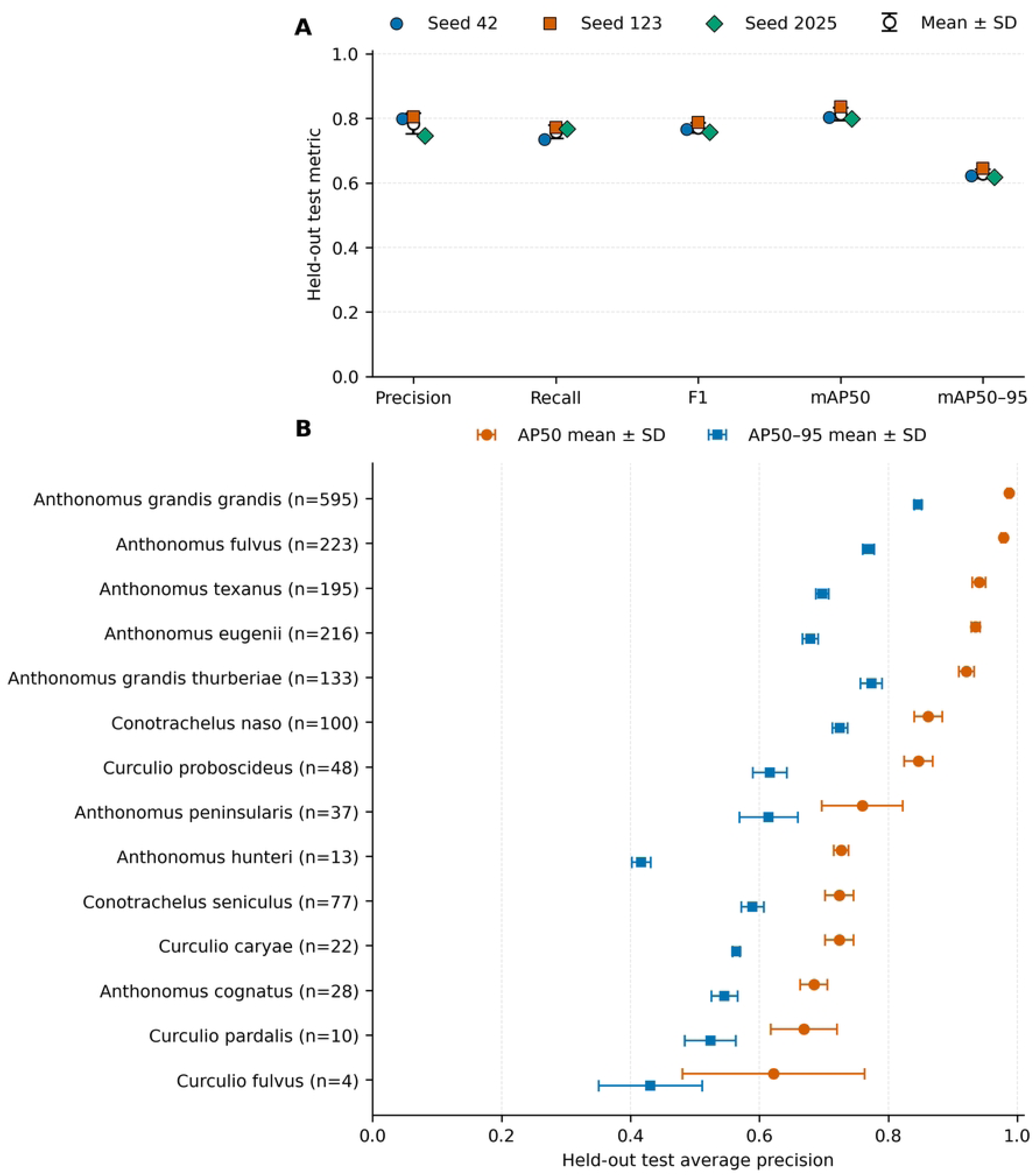
Held-out test performance of three variance-matched COCO-pretrained YOLO11n models. (A) Overall precision, recall, F1, mAP50, and mAP50-95 for seeds 42, 123, and 2025; open symbols and error bars indicate the arithmetic mean ± sample SD across the three seeds. (B) Species-level AP50 and AP50-95 summarized as mean ± sample SD across the same three seeds; labels report the number of test instances (n) for each taxon.

The direction of these matched results is consistent with agricultural transfer-learning studies showing benefits of pretrained representations when target-domain training data are limited [56,57] and with object-detection evidence that appropriate reuse of pretrained visual features can improve downstream detection [58]. For RRWIT, the controlled comparison indicates that COCO-pretrained initialization contributed to both higher mean held-out detection performance and lower between-seed variability relative to random initialization under the matched training protocol. The persistence of species-level performance differences, however, indicates that pretrained initialization did not eliminate taxon-specific detection challenges.

### 3.3 EigenGradCAM reveals object-size-dependent attribution localization and classification-dominated failure patterns

The complete 1,685-image single-object audit separated species assignment from spatial localization. RRWIT correctly classified and localized 1,527 images (90.6%); among the remaining images, 137 were species misclassifications, six were localization failures, and 15 had no detection. Thus, residual errors were dominated by fine-grained species assignment rather than inability to place a bounding box around the insect. The proportion correctly classified and localized was 90.4% for tiny objects, 93.6% for small, 93.4% for medium, and 83.6% for large objects. These RRWIT-specific size strata were defined by relative ground-truth box area (<1%, 1–<10%, 10–<50%, and ≥50% of image area, respectively), and the metric required both correct species assignment and localization. The lower percentage in the large-object stratum therefore should not be interpreted as evidence that large objects were intrinsically more difficult to detect; rather, prediction success in this seed-0 RRWIT audit did not vary monotonically with relative object area. EigenGradCAM attribution was evaluated on the fixed error-enriched subset of 258 detected images and summarized in Fig 8. Representative maps were selected objectively by median enrichment within each size stratum. Ground-truth-box enrichment was highest and most variable for tiny objects and approached the area-proportional reference of 1 as object area increased. Continuous object area was strongly and inversely associated with enrichment for both correct predictions (Spearman rho = -0.71, n = 115) and incorrect predictions (rho = -0.80, n = 143). Because enrichment divides attribution fraction by box-area fraction, large values for tiny objects should not be interpreted as stronger or more biologically meaningful attention.

**Fig 8.**
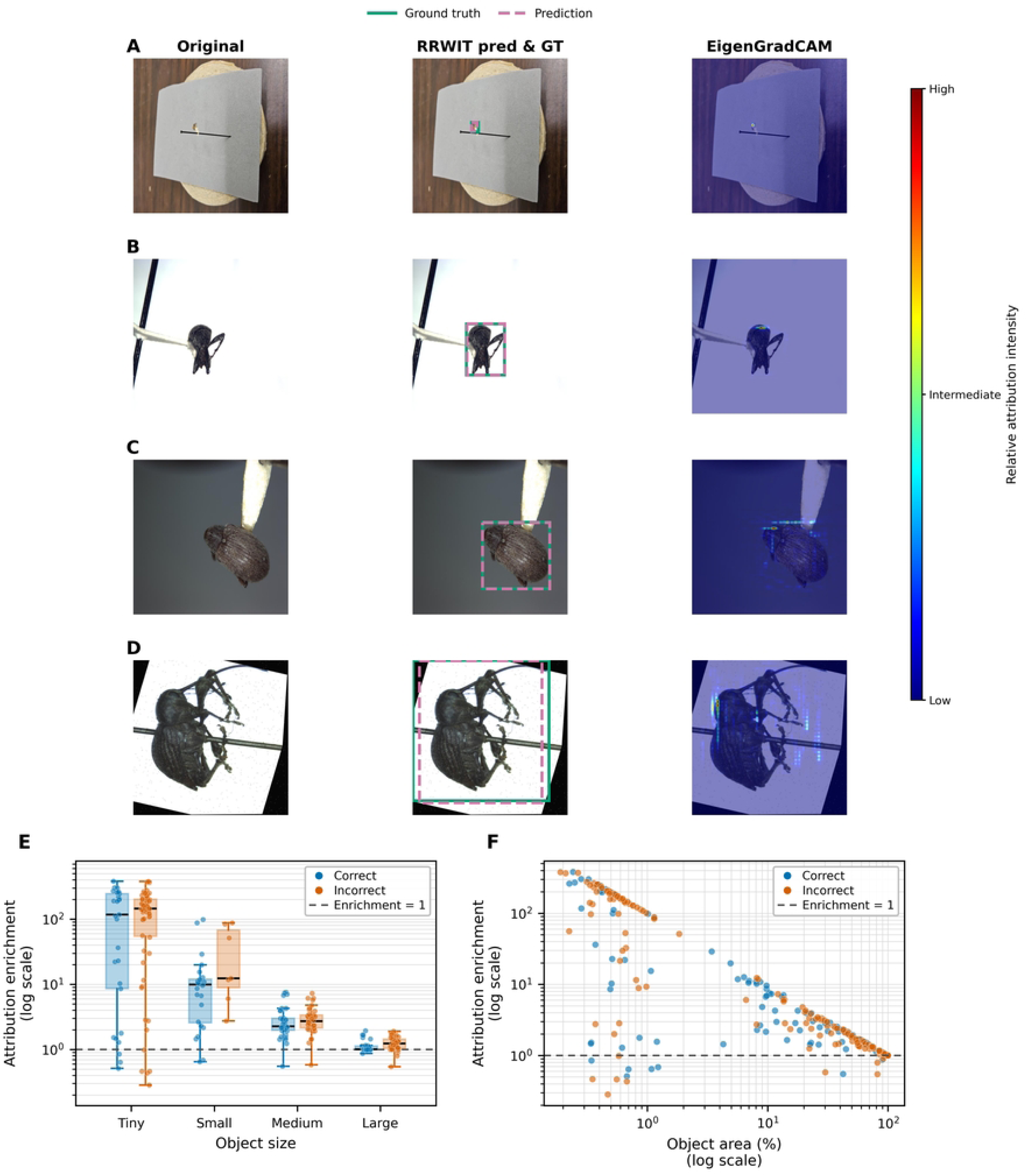
EigenGradCAM analysis of object-size-dependent attribution localization in RRWIT. (A-D) Objectively selected correctly detected examples for tiny, small, medium, and large ground-truth objects, showing the original image, prediction/ground truth, and class-targeted EigenGradCAM map from layer 16. (E) Ground-truth-box enrichment distributions for correct and incorrect detections in the 258-image error-enriched subset. (F) Continuous object area versus enrichment. Enrichment decreased with object area for correct (Spearman rho = -0.71, n = 115) and incorrect detections (rho = -0.80, n = 143). Attribution intensity is normalized within each image and is not an absolute cross-image scale.

Correct and incorrect predictions overlapped substantially in their localization patterns. An attribution map can therefore be concentrated on the insect even when the species label is wrong. This is expected for a fine-grained task in which a detector may identify the biologically relevant object yet fail to weight the specific visual characters needed for taxon discrimination. Grad-CAM-type methods are post hoc attribution tools, not causal explanations, and visual saliency can be misleading without quantitative controls [35,37]. The present analysis therefore supports a limited conclusion; RRWIT frequently derives class-associated evidence from the specimen region, but attribution localization alone does not establish taxonomically correct or biologically causal feature use.

The predominance of species-assignment errors explains why correct and incorrect detections can show similar object-focused attribution. The full audit contained recurrent substitutions such as *Conotrachelus seniculus* to *C. naso* and *Anthonomus peninsularis* to *A. texanus*, and the focal *Curculio* audit showed repeated confusion with *C. proboscideus*. These observations are consistent with the class-confusion structure in Fig 3 and with taxonomic literature describing subtle morphological distinctions within the *Anthonomus* and *Curculio* genera [49,50]. Collectively, these observations support diagnostic interpretation; RRWIT often localizes the relevant specimen correctly, but some residual failures occur at the finer species-discrimination stage rather than at gross object localization.

### 3.4 Targeted feature-scale and attention modifications do not outperform baseline RRWIT

Multi-scale feature fusion and attention are frequently used in agricultural detection when targets vary strongly in size or weak visual features must be amplified. Recent literature studies have likewise treated feature-fusion and attention modules as testable architectural hypotheses and evaluated them by ablation against an unchanged baseline [59,60]. We therefore used the object-size-dependent EigenGradCAM pattern to initiate, rather than predetermine, the RRWIT architecture screen.

The attribution analysis initiated a direct architectural hypothesis: additional shallow-scale information or explicit channel-spatial recalibration might improve performance if spatial feature preservation were a principal limitation. However, the seed-42 validation screen did not support that hypothesis. Baseline A0 achieved precision 0.801, recall 0.791, mAP50 0.842, and mAP50-95 0.648. A1 (P2-to-P3 assistance) fell to 0.672, 0.656, 0.718, and 0.536; A2 (P3 CBAM) reached 0.786, 0.762, 0.820, and 0.635; and A3 (combined) reached 0.814, 0.685, 0.787, and 0.611. A0 therefore remained best on the prespecified primary metric (S4 Fig A).

P2 assistance increased parameters from about 2.593 to 2.653 million and computation from 6.45 to 7.24 GFLOPs without improving accuracy, whereas P3 CBAM added negligible computational cost but still did not surpass A0. All four checkpoints passed FP32 ONNX export/runtime-parity checks, and the test partition remained unused during screening. Attention and multi-scale modules can improve other agricultural tasks [59,60], but their value is task- and placement-dependent. Here, the negative ablation result indicates that the residual RRWIT errors were not resolved by these three lightweight modifications and supports retaining the simpler baseline for the formal benchmark (S4 Fig B).

The P2-assisted pathway produced the clearest negative result. Relative to A0, A1 reduced validation mAP50-95 by 0.112 and mAP50 by 0.124 while increasing theoretical computation by about 12.2%. A2 was much closer to baseline, reducing mAP50-95 by only 0.013 with negligible computational overhead, showing that the result should not be generalized to attention mechanisms as a class. A3 illustrates why a single operating-point statistic should not determine model selection; its precision increased slightly to 0.814, but recall fell to 0.685 and both mAP metrics declined. The prespecified mAP50-95 criterion therefore retained A0 (S4 Figs A and B).

### 3.5 Cross-framework benchmarking confirms competitive RRWIT performance while newer YOLO architectures improve localization accuracy

The controlled benchmark compared eight detector configurations under identical source-image partitions and frozen validation-to-test procedures. The architectures, implementations, reference input sizes, and published model characteristics are summarized in S1 Table. This breadth was intended to compare RRWIT not only with preceding and subsequent YOLO generations but also with one-stage mobile, compound-scaled, and two-stage detector designs.

The validation-stage analysis established the framework ranking before locked-test evaluation (S5 Fig). Across seeds 42, 123, and 2025, YOLO26n achieved the highest validation mAP50– 95 (0.834 ± 0.006), followed by YOLO12n (0.819 ± 0.006) and RRWIT (0.808 ± 0.009). YOLOv8n, EfficientDet-D0, and YOLOv5nu followed at 0.790 ± 0.009, 0.776 ± 0.006, and 0.775 ± 0.007, respectively, whereas Faster R-CNN-320 and SSDLite320 produced substantially lower values of 0.673 ± 0.007 and 0.648 ± 0.005. Validation mAP50 was high for the YOLO-family models, reaching 0.988 ± 0.003 for YOLO26n and 0.971 ± 0.007 for RRWIT. Macro and support-weighted F1 scores (S5 Fig C) showed a similar high-performing group, although their ordering was not identical to mAP50–95 because these statistics reflect a fixed confidence operating point rather than integration over the precision-recall curve.

The validation analysis also revealed substantial dependence on object scale (S5 Fig D). Using the standard COCO object-size categories employed in the benchmark evaluation, AP for large objects was generally higher than for medium and especially small objects across frameworks, although the magnitude of this difference varied among architectures. This pattern is consistent with the well-established difficulty of detecting small objects, which contain fewer pixels and provide less spatial and feature information for localization and classification [15,17]. Importantly, this benchmark result is not directly comparable with the non-monotonic size pattern observed in the seed-0 RRWIT audit in Section 3.3. That analysis used RRWIT-specific relative-area categories, including a separate tiny-object class, and reported the proportion of images that were both correctly classified and localized for a single historical checkpoint, whereas S5 Fig D reports COCO-style size-stratified AP across independently trained benchmark models. YOLO26n showed particularly strong performance across the three COCO scale strata, supporting its validation-stage performance. The per-species comparison between RRWIT and YOLO26n further indicated that the aggregate advantage of YOLO26n was distributed across multiple taxa rather than being attributable to only one species (S5 Fig E, F).

Held-out test partitions subset results broadly preserved validation stage performance (Fig 9). YOLO26n had the highest mAP50-95, 0.810 ± 0.006, followed by YOLO12n at 0.804 ± 0.012 and RRWIT at 0.768 ± 0.004. RRWIT therefore ranked third of eight on the primary metric and remained close to EfficientDet-D0 (0.758 ± 0.007) and YOLOv8n (0.757 ± 0.010), while exceeding YOLOv5nu, Faster R-CNN-320, and SSDLite320 (Fig 9A Table 1).

**Fig 9.**
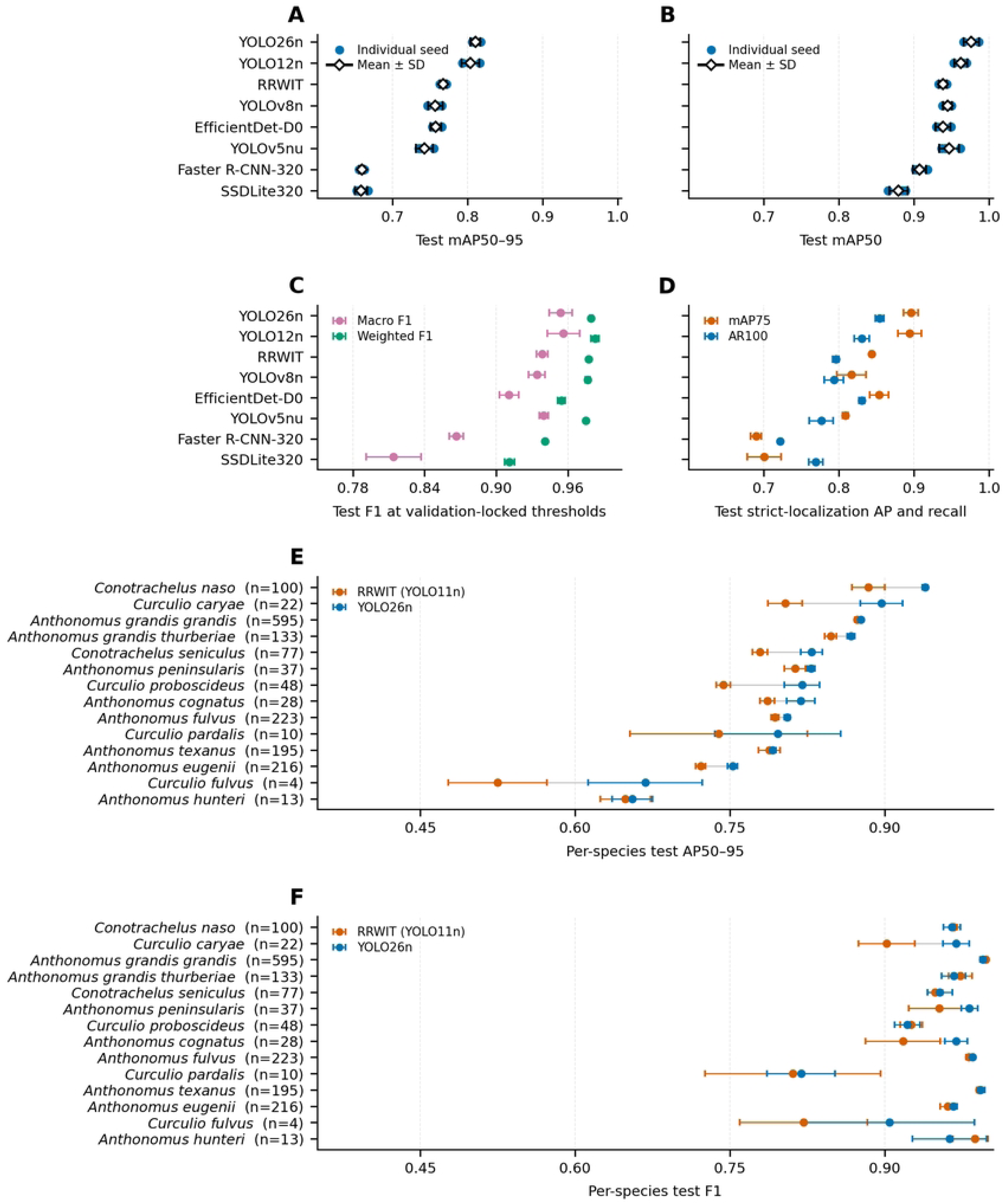
Held-out test partitions subset performance of eight object-detection frameworks under the frozen three-seed benchmark. **(A)** Test mAP50–95 and **(B)** test mAP50 across seeds 42, 123, and 2025. Blue circles indicate individual seed runs; open diamonds and horizontal error bars indicate arithmetic mean ± sample SD. **(C)** Macro and support-weighted F1 calculated using confidence thresholds selected exclusively on the validation partition and frozen before test evaluation. **(D)** Strict-localization performance summarized by mAP75 and average recall at a maximum of 100 detections per image (AR100). **(E)** Per-species test AP50–95 and **(F)** per-species test F1 comparing RRWIT (YOLO11n) with the highest-ranked framework, YOLO26n. Horizontal error bars in panels E and F represent sample SD across the three seeds; gray connecting lines aid within-species comparison, and *n* denotes the number of test annotations for each species.

**Table 1.**
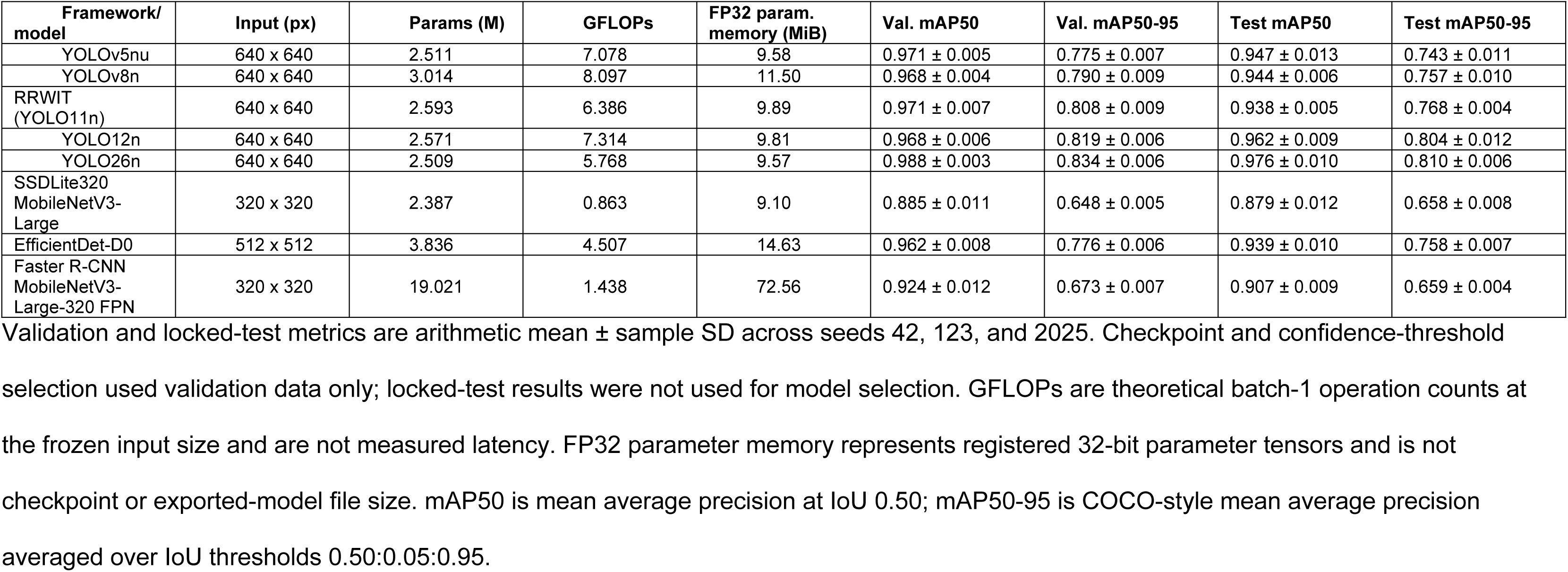
Predictive performance and computational footprint of the eight trained object-detection frameworks under the frozen WIT benchmark.

Metric dependence refined the comparison between models (Fig 9). RRWIT test mAP50 was 0.938 ± 0.005, compared with 0.976 ± 0.010 for YOLO26n and 0.962 ± 0.009 for YOLO12n; YOLOv5nu, YOLOv8n, and EfficientDet-D0 were numerically similar to RRWIT at IoU 0.50 despite lower mAP50-95 (Fig 9B). At IoU 0.75, RRWIT reached 0.844 ± 0.002 versus 0.896 ± 0.010 for YOLO26n and 0.894 ± 0.016 for YOLO12n (Fig 9D). This pattern indicates that the newer YOLO architectures gained primarily under stricter localization requirements. RRWIT nevertheless retained high support-weighted F1 (0.977 ± 0.001) and low between-seed mAP50-95 SD (0.004) (Fig 9C).

Average recall at a maximum of 100 detections per image (AR100) showed a related pattern: YOLO26n reached 0.854 ± 0.006, YOLO12n and EfficientDet-D0 approximately 0.830, and RRWIT 0.796 ± 0.004 (Fig 9D). At validation-selected confidence thresholds, macro F1 was 0.956 ± 0.014 for YOLO12n, 0.954 ± 0.010 for YOLO26n, and 0.939 ± 0.005 for RRWIT (Fig 9C). These operating-point F1 values and COCO-style AP are complementary rather than interchangeable: F1 summarizes one frozen threshold, whereas AP integrates ranked predictions and, for mAP50-95, multiple localization thresholds.

Validation-to-test changes also varied across frameworks. RRWIT changed from 0.808 ± 0.009 validation mAP50-95 to 0.768 ± 0.004 on locked test (approximately -0.040), compared with changes of approximately -0.024 for YOLO26n and -0.015 for YOLO12n (Table 1). The broad ranking nevertheless remained stable, providing a useful internal generalization check.

The benchmark also places the RRWIT model choice in a temporal context. YOLO11n was a compact current-generation detector when RRWIT development was designed, whereas YOLO12 (attention-centric real-time detector design) and YOLO26 (introduced a newer architecture incorporating a lighter regression head and native end-to-end inference capabilities) became available later [22,41,42]. Similar within-dataset agricultural comparisons increasingly showed that the model with the highest accuracy is not necessarily the model with the preferred deployment trade-off [61,62]. In the present dataset, YOLO26n improved strict-IoU accuracy while using slightly fewer parameters and lower theoretical computation than RRWIT, making it the strongest evaluated candidate for future RRWIT development without implying that its ranking will remain unchanged as our biological dataset and computer vision architectures evolves. The present benchmark therefore should not be interpreted as evidence that the initial YOLO11n choice was inappropriate; rather, it demonstrates how the performance frontier changed as newer architectures became available.

### 3.6 Species-level performance identifies persistent challenges among visually similar and underrepresented taxa

Species-level evaluation revealed substantially greater heterogeneity than was apparent from framework-level summary metrics. Across the three prespecified seeds, RRWIT showed strong locked-test performance for several taxa, including *Conotrachelus naso* and *Anthonomus grandis grandis*, whereas lower AP50–95 occurred for a subset of taxa including *Curculio fulvus*, *Anthonomus hunteri*, *Anthonomus eugenii*, and *Curculio pardalis*. Importantly, test representation varied markedly among species, from 595 annotated *A. grandis grandis* and 223 *A. fulvus* objects to only 22 *C. caryae*, 13 *A. hunteri*, 10 *C. pardalis*, and four *C. fulvus*. Consequently, the larger between-seed uncertainty observed for some sparsely represented taxa, particularly *C. fulvus* and *C. pardalis*, should be interpreted in the context of their limited test support rather than as precise estimates of population-level performance (Fig 10A).

**Fig 10.**
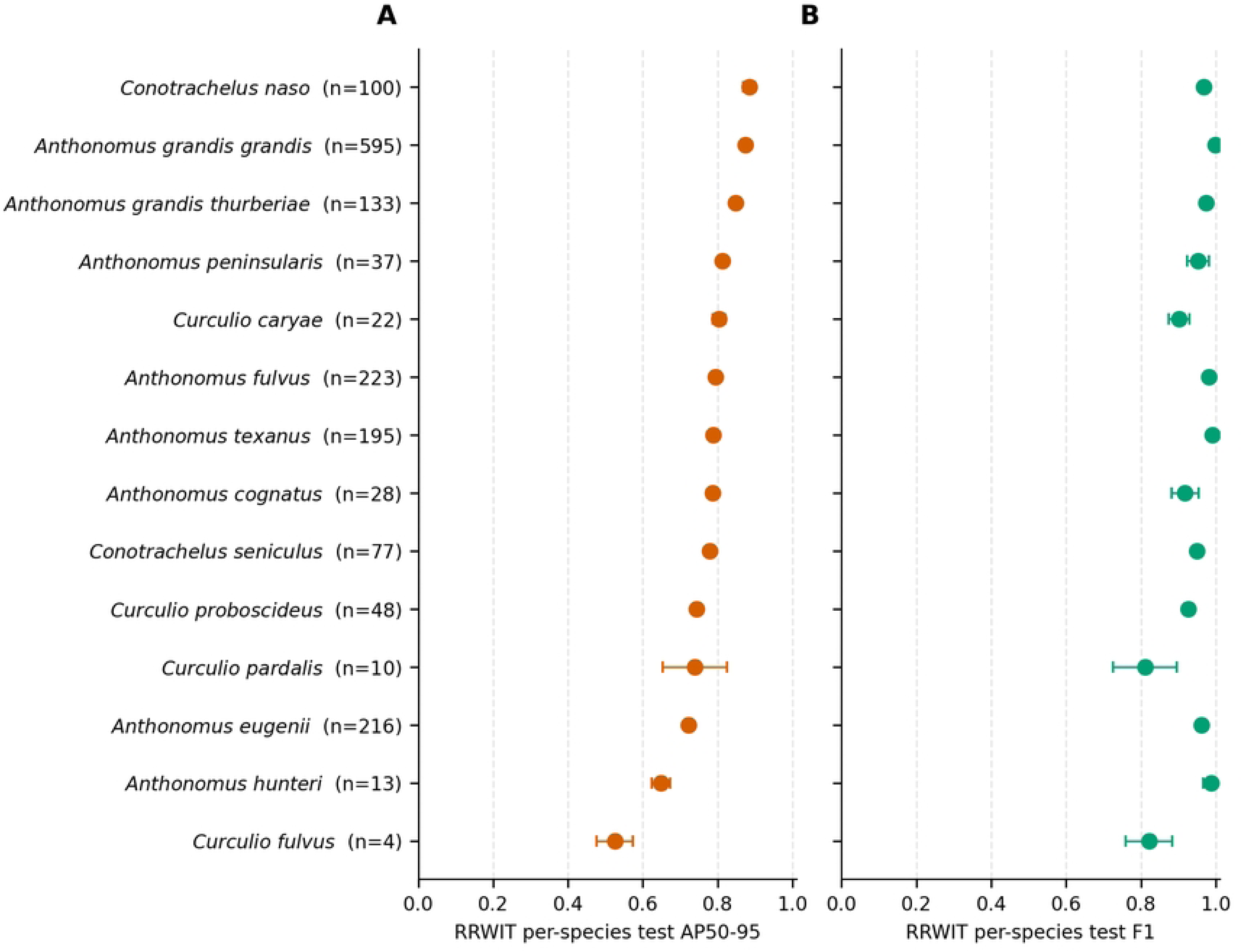
Species-level locked-test performance of RRWIT across three independent training seeds. (A) Species-specific AP50-95 and (B) F1 at validation-selected confidence thresholds. Points are arithmetic means and horizontal error bars are sample SD; numbers in parentheses on y-axis show locked-test annotation support.

AP50–95 (Fig 10A) and F1 (Fig 10B) also provided complementary information about the origin of species-level errors. Some classes with comparatively low AP50–95 nevertheless maintained high F1 scores at the validation-selected operating threshold (Fig 10). This was particularly evident for *A. hunteri*, indicating that successful detection and class assignment at IoU 0.50 can coexist with reduced localization consistency when performance is averaged over progressively stricter IoU thresholds from 0.50 to 0.95. Conversely, *C. fulvus* and *C. pardalis* showed reductions in both AP50–95 and F1, although interpretation is necessarily constrained by their very small test supports. Thus, low species-level AP should not automatically be interpreted as a pure classification failure; it can reflect some combination of localization precision, missed detections, species substitution, and threshold-dependent classification performance.

The three-seed confusion matrices reinforced this pattern (Figs 11, 12 and S6) with most classes remaining strongly diagonal. The mean on the held-out previously unseen test partitions subset diagonal proportions were 1.00 for *A. grandis grandis*, 0.99 for *A. peninsularis* and *A. texanus*, 0.98 for *A. fulvus* and *C. naso*, and 0.95 for *A. eugenii*. *Curculio* taxa with lower number of images were more heterogeneous, with corresponding means of approximately 0.91 for *C. caryae*, 0.75 for *C. fulvus*, 0.87 for *C. pardalis*, and 0.90 for *C. proboscideus*, together with more visible substitutions and unmatched-ground-truth proportions (Fig 11). Fine-grained *Anthonomus* and *Curculio* identification depends on combinations of subtle morphological characters [49,50], providing a biological reason why some distinctions are difficult in two-dimensional images. Class abundance was not a complete explanation; however, some taxa with lower number of images performed well, and some more represented taxa remained harder than the best classes. Long-tailed support and high inter-/intra-class visual variation are recognized challenges in insect computer vision [63–65].

**Fig 11.**
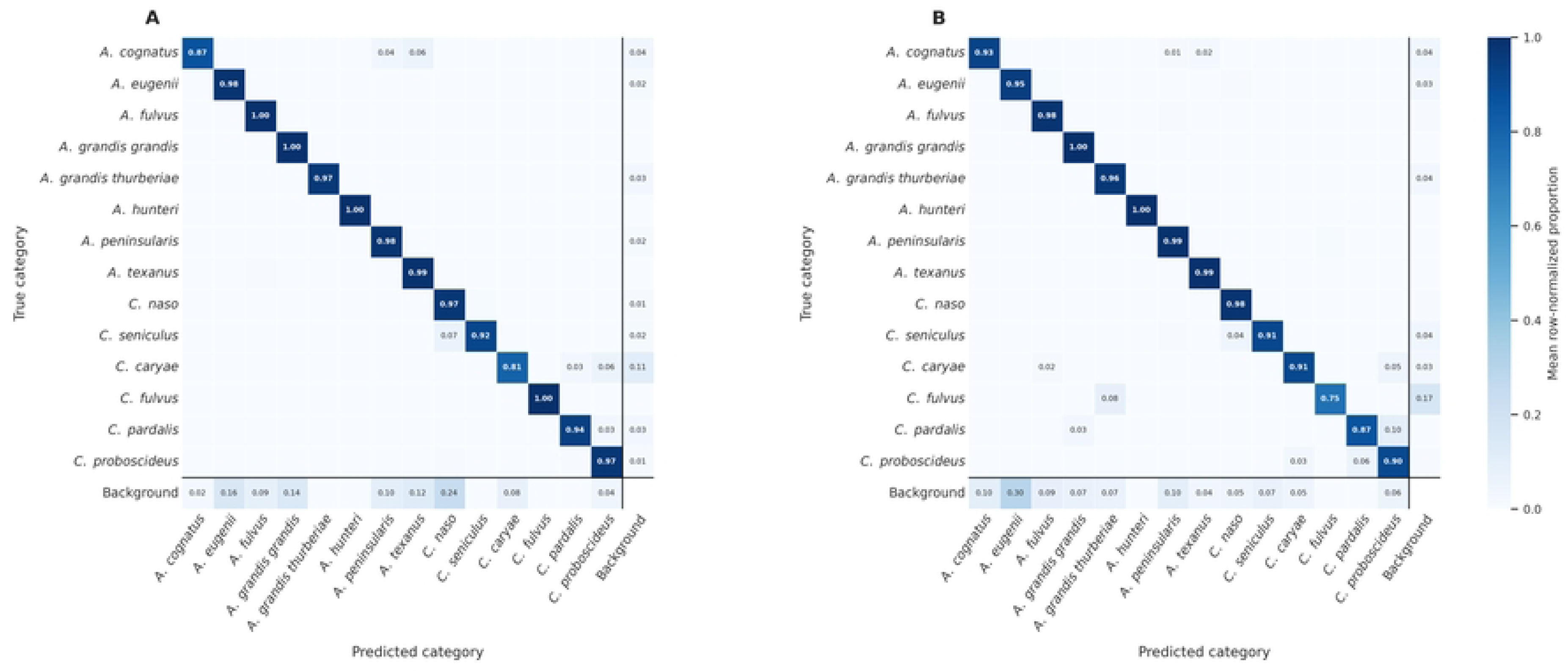
Validation and held-out test partitions subset confusion patterns for RRWIT (YOLO11n) across three independent training seeds. Mean row-normalized confusion matrices are shown for the validation set (A) and locked test set (B). Each cell is the arithmetic mean across seeds 42, 123, and 2025 after independent row normalization. Rows are true categories and columns are predicted categories. Off-diagonal biological cells indicate species substitutions, the Background column represents unmatched ground-truth objects (false negatives), and the Background row represents unmatched predictions (false positives) by predicted class. The Background-Background cell is undefined and left blank. Matching and thresholding followed the frozen evaluation procedure.

The difficult classes were not unique to RRWIT. YOLO26n showed a broadly similar taxonomic error structure (Fig 12), and YOLO12n showed the same persistent difficulty for several low-support *Curculio* taxa (S6 Fig). For YOLO26n, the mean held-out previously unseen test partitions subset diagonal proportions for *C. fulvus*, *C. pardalis*, and *C. proboscideus* were approximately 0.83, 0.83, and 0.90, respectively. YOLO12n showed approximately 0.83, 0.90, and 0.95 for the same taxa (Fig 12; S6 Fig). Cross-framework persistence suggests that a substantial component of residual error reflects task/data properties, limited representation, subtle morphology, orientation, and visibility of diagnostic structures rather than a single RRWIT architecture. Related insect-monitoring studies similarly emphasize field-representative retraining and the difficulty of species-level discrimination among morphologically uniform groups [20,66].

**Fig 12.**
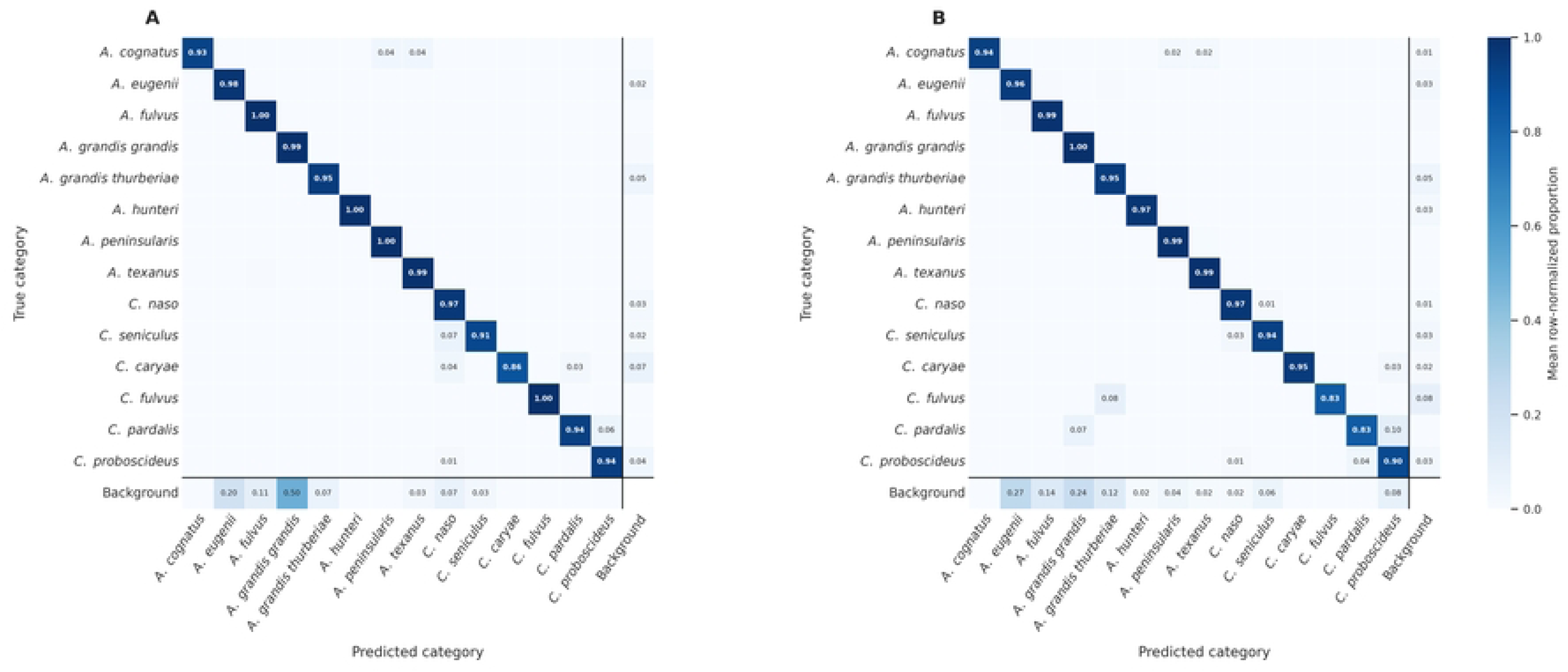
Validation and held-out test partitions subset confusion patterns for YOLO26n across three independent seeds. Mean row-normalized confusion matrices are shown for the validation set (A) and locked test set (B). Each cell is the arithmetic mean across seeds 42, 123, and 2025 after independent row normalization. Rows are true categories and columns are predicted categories. Off-diagonal biological cells indicate species substitutions, the Background column represents unmatched ground-truth objects (false negatives), and the Background row represents unmatched predictions (false positives) by predicted class. The Background-Background cell is undefined and left blank. Matching and thresholding followed the frozen evaluation procedure.

The combination of Fig 10 and the multi-framework confusion matrices indicates that future improvements should prioritize targeted acquisition of diverse images for difficult taxa, especially low-support *Curculio* species, while deliberately varying specimen orientation, scale, illumination, background, sex where relevant, and visibility of diagnostic morphological structures. Such targeted enrichment would be more informative than indiscriminately increasing already well-represented classes. From an applied surveillance perspective, these species-level results also define where human verification remains most valuable. Automated inference can substantially reduce the burden of screening large numbers of trap images, but predictions involving low-support or repeatedly confused taxa should be treated with greater caution than predictions for consistently high-performing classes. This is particularly important when species identity would trigger a management or regulatory response. Accordingly, RRWIT is better viewed as a decision-support system for rapid taxonomic screening than as a replacement for expert confirmation in the most difficult cases. The three-seed analyses make that limitation explicit while also showing that the majority of species are detected with highly consistent class-level performance.

### 3.7 Accuracy-complexity trade-offs favor lightweight one-stage detectors

The leading models occupied a narrow lightweight regime rather than obtaining higher accuracy through substantially greater capacity (Table 1). YOLO26n combined test mAP50-95 of 0.810 ± 0.006 with 2.509 million parameters, 5.768 GFLOPs, and 9.57 MiB of FP32 parameter memory; YOLO12n used 2.571 million parameters and 7.314 GFLOPs; RRWIT used 2.593 million and 6.386 GFLOPs. By contrast, SSDLite320 had only 0.863 GFLOPs but lower mAP50-95 (0.658 ± 0.008), while Faster R-CNN-320 contained 19.021 million parameters and 72.56 MiB of FP32 parameter memory yet reached 0.659 ± 0.004.

These observations illustrate why parameter count and theoretical operation count should not be interpreted as runtime. The models also used different frozen input sizes: the YOLO-family detectors used 640 × 640 pixels, EfficientDet-D0 512 × 512, and the SSDLite/Faster R-CNN configurations 320 × 320; therefore, actual speed depends on operator structure, memory access, inference-engine implementation, preprocessing, post-processing, and hardware utilization [67]. Recent agricultural work similarly treats accuracy, model size, and computation as joint design objectives rather than optimizing a single scalar [68–70]. For this weevil task, compact YOLO-family detectors occupied the most favorable accuracy-complexity region among the evaluated configurations; actual deployment performance required direct runtime measurement.

Within these constraints, the strongest result is not that all one-stage detectors are universally superior, but that the compact YOLO-family models occupied the most favorable accuracy-complexity region for this particular fine-grained task. Similar parameter budgets did not yield equivalent localization performance, and YOLO26n improved the RRWIT accuracy-complexity frontier without moving to a substantially larger network (Table 1). This makes lightweight one-stage detection a defensible design space for future RRWIT development while leaving actual device performance to direct deployment measurements.

### 3.8 INT8 quantization substantially reduces model size but does not guarantee faster CPU inference

Common static INT8 quantization substantially compressed all four deployment candidates, but accuracy retention was architecture-dependent (Table 2). YOLO26n, YOLO12n, and RRWIT retained 98.6%, 98.8%, and 98.1% of FP32 validation mAP50-95, respectively, while their serialized ONNX artifacts shrank by approximately 69-71%. RRWIT decreased from 10.09 to 2.91 MiB (71.2%). Under the prespecified aggregate gate, only YOLO12n met all three accuracy tolerances; YOLO26n and RRWIT exceeded the mAP50-95 loss threshold slightly while retaining mAP50 and weighted F1 within tolerance. This is consistent with the storage benefit expected from lower-precision integer representations [71].

**Table 2.**
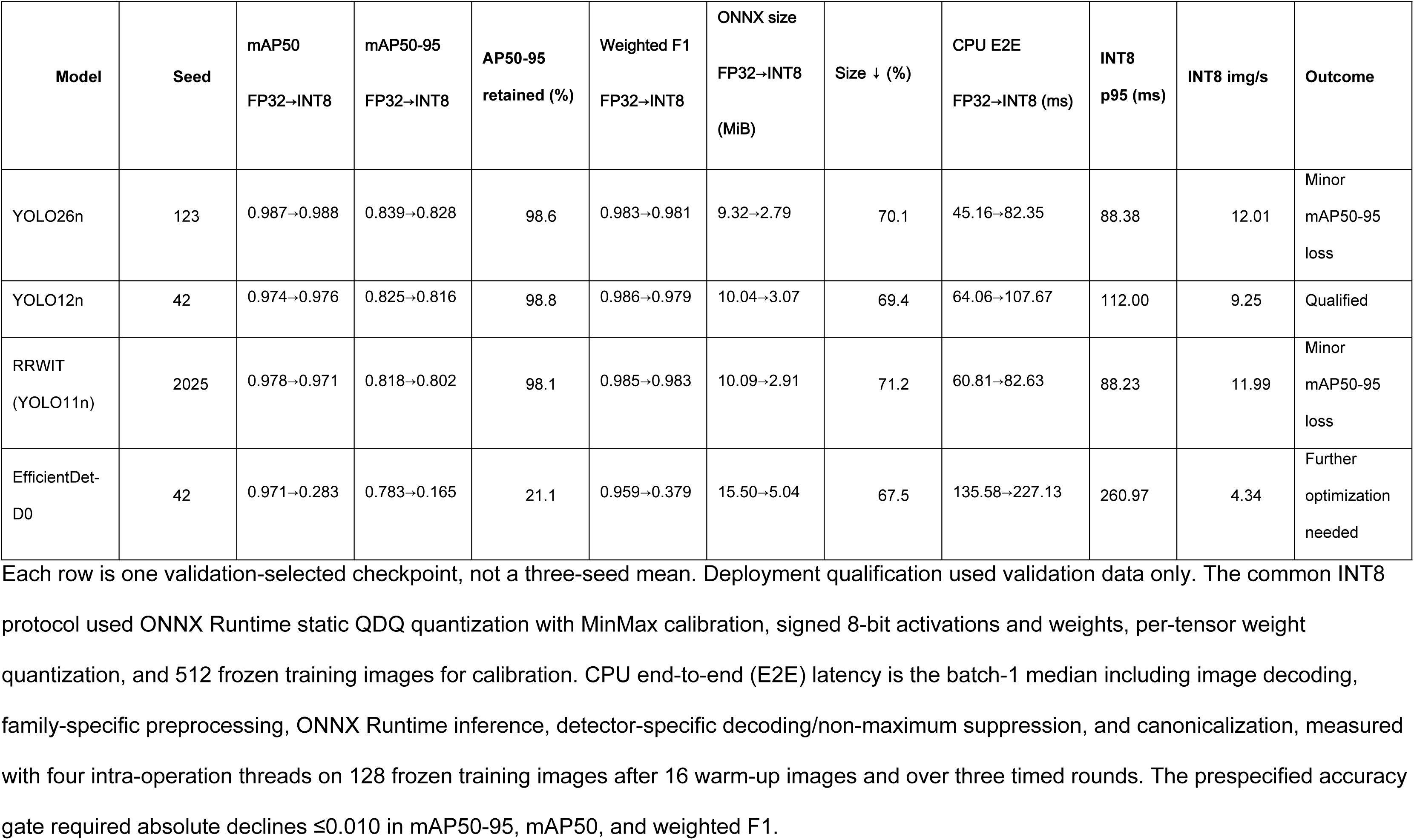
Validation accuracy, serialized ONNX model size, and CPU inference characteristics of four validation-selected deployment candidates after common INT8 post-training quantization.

Compression did not accelerate the tested CPU execution path. Median end-to-end latency increased from 45.16 to 82.35 ms/image for YOLO26n, 64.06 to 107.67 for YOLO12n, 60.81 to 82.63 for RRWIT, and 135.58 to 227.13 for EfficientDet-D0. Integer inference can be faster on software/hardware stacks optimized for low-precision kernels, but quantization format alone does not guarantee speed. Hardware-targeted deployment studies illustrate that device-specific runtimes can realize acceleration not observed under a generic CPU path [72,73].

This result is compatible with hardware-targeted quantization studies rather than contradictory to them. Wang et al. converted YOLOv11 to an INT8 RKNN representation optimized for an RK3568 edge device and reported a size reduction from 10,156 to 4,194 kB together with a single-image detection-time reduction from 52.1 to 33.6 ms [72]. Chen et al. likewise evaluated a YOLO11-derived model on a Jetson TX2 under INT8 CPU-only deployment and reported strong accuracy retention [73]. Their results illustrate the benefit when graph conversion, integer kernels, and hardware are co-optimized; our experiment instead tested a common ONNX Runtime CPU route across architectures.

EfficientDet-D0 was particularly sensitive to the common quantization protocol. For example, mAP50-95 fell from 0.783 to 0.165 and weighted F1 from 0.959 to 0.379 despite a 67.5% size reduction (Tables 2 and 3). Per-channel quantization substantially recovered accuracy, with U8U8/per-channel reaching mAP50-95 of 0.556 and weighted F1 of 0.800, but no diagnostic configuration restored FP32-equivalent accuracy or latency. A bridge-control analysis reproduced the calibration bytes, preprocessing stream, quantization parameters, and common artifacts, excluding artifact drift as an explanation for the collapse. Thus, quantization should be treated as an architecture- and hardware-specific optimization problem rather than a universal conversion step.

**Table 3.** Architecture-specific EfficientDet-D0 INT8 quantization diagnostics following substantial accuracy degradation under the common quantization protocol used in Table 2.

| **Variant** | **Role** | **mAP50** | **mAP50-95** | **Weighted F1** | **ΔmAP50-95 vs FP32** | **ONNX size (MiB)** | **CPU E2E median (ms)** | **Throughput (img/s)** |
|---|---|---|---|---|---|---|---|---|
| FP32 | Reference | 0.971 | 0.783 | 0.959 | 0.000 | 15.50 | 135.58 | 7.36 |
| S8S8/PT | Common protocol | 0.283 | 0.165 | 0.379 | −0.618 | 5.04 | 227.13 | 4.34 |
| S8S8/PC | Diagnostic only | 0.768 | 0.544 | 0.789 | −0.238 | 5.47 | 205.78 | 4.79 |
| U8U8/PT | Diagnostic only | 0.415 | 0.266 | 0.526 | −0.517 | 5.04 | 182.32 | 5.38 |
| U8U8/PC | Diagnostic only | 0.785 | 0.556 | 0.800 | −0.227 | 5.47 | 166.48 | 5.94 |

FP32 is the reference and S8S8/PT is the authoritative common cross-framework INT8 protocol. S8S8/PC, U8U8/PT, and U8U8/PC were architecture-specific diagnostics only. S8S8 and U8U8 denote signed and unsigned 8-bit activation/weight formats; PT and PC denote per-tensor and per-channel weight quantization. Validation metrics used the same 1,690-image validation set; the locked test set remained prohibited. CPU E2E timing followed the controlled batch-1 ONNX Runtime protocol used for Table 2.

Deployment experiments therefore separate four properties (theoretical computation, serialized storage, predictive retention after quantization, and measured runtime) that are often conflated. RRWIT can be compressed to a roughly 3-MiB INT8 artifact while retaining high validation accuracy, but speed claims require backend- and device-specific testing. For RRWIT specifically, INT8 therefore provides a viable route to substantially lower serialized storage, but additional backend- and hardware-specific optimization is required before claiming a speed advantage. The current cloud-hosted prototype supports both image upload and live capture and returns bounding boxes, species labels, and confidence scores through an interactive interface (S1 Video). It currently operates in a CPU-based cloud environment (2 vCPUs and 16 GB RAM). Application-level latency and scalability were not systematically benchmarked, so apparent web-response time is not reported as a deployment metric. Other metrics such as energy use, and representative mobile/edge performance also remain to be evaluated.

### 3.9 Implications, limitations, and future development of RRWIT

Taken together, the experiments support the original design premise without implying that YOLO11n is universally optimal. Pretrained initialization improved mean performance and reduced between-seed variability. Quantitative EigenGradCAM showed that attribution localization is strongly scale-dependent and that object-focused attribution does not ensure correct species classification. Furthermore, the targeted A1-A3 modifications failed to improve the baseline and the frozen benchmark showed that newer YOLO12n and especially YOLO26n can improve strict-IoU localization at a comparable lightweight footprint. RRWIT is therefore best understood as an evolving identification framework, with YOLO26n representing a strong candidate backbone for future versions rather than a guaranteed permanent successor.

Confusion patterns further showed that the remaining classification errors were not distributed uniformly across taxa (Fig 11). RRWIT retained particularly strong discrimination of the boll weevil, *A. grandis grandis*, for which the mean diagonal proportion across the three independently trained models was 99.66% on the locked test set. Residual substitutions from this pest species were rare and individually below the 2% annotation threshold used in Fig 11. In contrast, the pecan weevil, *C. caryae*, remained more challenging: its mean test diagonal was 90.91%, with 4.55% assigned to *C. proboscideus*, 1.52% to *A. fulvus*, and 3.03% unmatched as background. Importantly, most matched wrong-species assignments across the test evaluations occurred between non-pest taxa rather than across the pest/non-pest boundary, indicating that a substantial component of the remaining species-level error involved discrimination among morphologically similar non-target weevils rather than systematic confusion of the two target pest species (Fig 11).

The newer YOLO26n architecture reduced several of these residual errors while preserving the same general pattern (Fig 12). Its mean locked-test diagonal proportions reached 99.72% for *A. grandis grandis* and 95.45% for *C. caryae*. For pecan weevil, the principal residual substitution was again *C. proboscideus* (3.03%), with 1.52% assigned to background. Across the three test evaluations, the great majority of YOLO26n matched wrong-species assignments likewise occurred among non-pest taxa. Thus, the confusion analysis identifies two complementary features of the benchmark: boll-weevil discrimination was highly stable across both detectors, whereas the more persistent errors were concentrated among particular morphologically similar taxa, especially within the more difficult *Curculio* group (Figs 11 and 12). These results support targeted expansion of underrepresented and difficult species in future RRWIT datasets rather than relying solely on architectural improvements to resolve the remaining taxonomic ambiguity.

The current internal test partition preserves image sources and historical acquisition contexts represented in the overall dataset. It is not intended to serve as external validation from new locations, institutions, trapping programs, devices, dates, or end users. Insect-monitoring research emphasizes that incomplete, imbalanced reference libraries and changing environmental context can drive errors, including assignment of unfamiliar taxa to visually similar known classes [74]. Future RRWIT upgrades will therefore prioritize improving annotation from bounding boxes to instance segmentation, keypoints detections, and Panoptic segmentation. It will also include diverse new images of low-support and repeatedly confused taxa, independent external testing, and mechanisms for flagging unfamiliar or uncertain specimens rather than assuming every insect belongs to one of the 14 trained classes. An expert-in-the-loop cycle in which such specimens are reviewed, annotated, and incorporated into later training releases would allow the biological reference library to expand while retaining taxonomic oversight [74].

As our image collection expands, the scientific goals of RRWIT can move beyond simply localizing a weevil and identifying its taxon toward investigating the specific morphological structures that inform those predictions. Future enhancements could incorporate instance segmentation to isolate complete insect silhouettes and anatomical regions, and utilize taxonomist-defined keypoints to encode homologous landmarks relevant to diagnostic features. Mask R-CNN introduced a unified framework for object detection and instance segmentation and can be extended to estimate keypoints [75], while DeepPoseKit demonstrated fast and robust dense keypoint estimation for insects and other animals [76]. Integrating these richer annotation types would enable direct, biologically meaningful comparisons between expert-defined taxonomic characters and attribution maps derived using techniques such as EigenGradCAM particularly in complex images containing multiple insects, debris, vegetation, and trap components.

Whole-trap or field imagery may eventually justify scene-level methods. Panoptic segmentation unifies individual-object and semantic scene interpretation [77], and PhenoBench demonstrates coordinated semantic, instance, and hierarchical agricultural image tasks, including evaluation on an unseen field [78]. In parallel, direct benchmarking on intended ARM CPUs, neural-processing units, or other accelerators is required before on-device performance can be claimed.

The prototype also contains an exploratory conversational artificial intelligence (AI) assistant (S1 Video), illustrating a longer-term path toward combining visual identification with curated taxonomic, biological, surveillance, and management information. That component was not systematically evaluated and is not part of the detection benchmark. Any future decision-support implementation would require separate assessment of retrieval accuracy, factual reliability, uncertainty communication, provenance, usability, and safeguards against unsupported responses. The current study therefore establishes a technically evaluated visual-identification foundation while keeping these broader capabilities explicitly prospective.

## 4 Conclusion

RRWIT demonstrates that transfer learning with a lightweight detector can support accurate identification of morphologically similar pests and non-pest weevils. Replicated initialization controls, quantitative EigenGradCAM, targeted ablation, and the frozen eight-framework comparison provide a substantially stronger basis than a single-model evaluation. RRWIT remained competitive, but YOLO26n achieved higher strict-IoU localization accuracy with a comparable or smaller computational footprint (Table 1), showing that RRWIT should be treated as an extensible framework rather than a system permanently tied to YOLO11n.

The results also identify two priorities for future development. First, species-level errors and low support for several taxa make representative biological data and independent external validation central to further gains. Second, model compactness is not equivalent to execution speed. INT8 quantization reduced RRWIT from 10.09 to 2.91 MiB but did not improve end-to-end computational latency on the tested CPU/ONNX Runtime path (Table 2). Deployment claims should therefore be based on measurements from the intended hardware and runtime.

With continued expert-verified image acquisition, external validation, and hardware-specific optimization, RRWIT provides a practical foundation for progressively richer automated weevil identification. Future work can evaluate newer lightweight backbones and, as data permit, instance segmentation, anatomically defined keypoints, and complex-scene interpretation while retaining expert oversight for uncertain or management-critical cases. The existing web prototype demonstrates an accessible interaction pathway, but operational field deployment and the exploratory conversational component require separate validation.

## Acknowledgements

We thank the Texas Boll Weevil Eradication Foundation, Texas Department of Agriculture, and the Texas Pecan Growers Association for field support and access to traps. We also thank the Texas A&M University Insect Collection for providing reference specimens. Special thanks to Jayda Arriaga and Kiley Stout for help with imaging and insect pinning. We would like to thank the ARS SCINet Office in general, and specifically Drs. Brian Stucky and Heather Savoy, for providing general computer vision training for our team as well as Joey Jones for his efforts in allocating GPU resources when they were most needed. Mention of trade names, companies, or commercial products in this publication is solely for the purpose of providing specific information and does not imply recommendation or endorsement of the products by the U.S. Department of Agriculture. The USDA is an equal opportunity provider and employer.

## Supporting Information

**S1 Fig. Bar plot showing the distribution of labeled images across the training, validation, and test subsets for each of the 14 weevil species.** The x-axis represents the species names, while the y-axis indicates the number of labeled images per species. Orange bars correspond to the training set, green bars to the validation set, and blue bars to the test set. A stratified sampling approach was used to ensure that all subsets retained proportional representation of each species.

**S2 Fig. Class distribution, spatial distribution, and bounding-box geometry of the annotated RRWIT dataset.** (A) Class frequencies, normalized bounding boxes, object-center distribution, and width-height distribution. (B) Univariate and pairwise distributions of normalized x-center, y-center, width, and height. These are dataset-level quality-control diagnostics and do not independently establish annotation correctness or biological independence among partitions.

**S3 Fig. Example output from the RRWIT for individual images.** Uploaded image showing automated detection and species classification with bounding box and predicted label confidence.

**S4 Fig. Validation performance and computational cost of targeted RRWIT architecture modifications.** (A) Seed-42 validation precision, recall, mAP50, and mAP50–95 for the unmodified RRWIT architecture (A0), P2-to-P3 feature assistance (A1), P3 Convolutional Block Attention Module refinement (A2), and the combined P2-assistance plus CBAM architecture (A3). A0 represents the unmodified RRWIT/YOLO11n architecture trained as the seed-42 matched transfer-learning control and is distinct from the historical seed-0 RRWIT.v1.0.0 checkpoint. (B) Parameter count and theoretical computational complexity at an input resolution of 640 × 640 pixels. Blue circles denote parameters and correspond to the left y-axis; orange squares denote GFLOPs and correspond to the right y-axis. Lines connect values only to aid visual comparison among the four discrete architectures. A0 achieved the highest prespecified primary validation metric, mAP50–95 (0.648). A2 incurred negligible additional computational cost but did not exceed A0, whereas the P2-containing A1 and A3 architectures increased computational demand without improving mAP50–95. All architectures were screened using the same seed and validation protocol; the held-out test partition was not used for architecture selection. Abbreviations: CBAM, Convolutional Block Attention Module; GFLOPs, billions of floating-point operations; mAP50, mean average precision at intersection over union 0.50; mAP50-95, mean average precision averaged over intersection-over-union thresholds 0.50 to 0.95.

**S5 Fig. Validation-stage performance of eight object-detection frameworks before locked-test evaluation.** (A) Validation mAP50–95 and (B) validation mAP50 across seeds 42, 123, and 2025. Blue circles represent individual seed runs; open diamonds and horizontal error bars represent arithmetic mean ± sample SD. (C) Macro and support-weighted F1 calculated at confidence thresholds selected on validation data. (D) Validation AP stratified by COCO object-size categories for small, medium, and large objects. (E) Per-species validation AP50–95 and (F) per-species validation AP50 comparing RRWIT (YOLO11n) with YOLO26n; horizontal error bars represent sample SD across the three seeds, and connecting lines aid within-species comparison. Overall class-averaged values are reported as mAP, whereas species-specific values are reported as AP. Abbreviations: AP, average precision; COCO, Common Objects in Context; F1, harmonic mean of precision and recall; mAP, mean average precision; SD, sample standard deviation.

**S6 Fig. Validation and locked-test confusion patterns for YOLO12n across three independent training seeds.** Mean row-normalized confusion matrices for YOLO12n on the validation set (A) and locked test set (B). Each cell represents the arithmetic mean across seeds 42, 123, and 2025 after independent row normalization. Rows are true categories and columns are predicted categories. Off-diagonal biological cells indicate species substitutions, the Background column represents unmatched ground-truth objects (false negatives), and the Background row represents unmatched predictions (false positives) by predicted class. The Background–Background cell is undefined and left blank. Matching and thresholding were identical to the frozen evaluation procedure used for the other framework-specific confusion matrices. Abbreviations: F1, harmonic mean of precision and recall; SD, sample standard deviation.

**S1 Table.** Published architecture and implementation characteristics of the eight object-detection frameworks evaluated in this study. Parameters and GFLOPs are published specifications for the corresponding reference/pretrained detector variants at the stated reference input size and are not measurements of the 14-class models fine-tuned in this study. Software license refers to the implementation used and does not supersede separate terms that may apply to pretrained weights, training datasets, or downstream deployment. **S1_Table.** Published architecture and implementation characteristics of the eight object-detection frameworks evaluated in this study.

| **Model** | **Detector architecture** | **Architecture reference** | **Implementation used** | **Reference input size (pixels)** | **Parameters (M)†** | **GFLOPs†** | **Software license‡** |
|---|---|---|---|---|---|---|---|
| YOLOv5nu | One-stage, anchor-free YOLO | [39] | Ultralytics 8.4.118 | 640 | 2.6 | 7.7 | AGPL-3.0 / Enterprise |
| YOLOv8n | One-stage, anchor-free YOLO | [40] | Ultralytics 8.4.118 | 640 | 3.2 | 8.7 | AGPL-3.0 / Enterprise |
| RRWIT (YOLO11n) | One-stage, anchor-free YOLO | [22] | Ultralytics 8.4.118 | 640 | 2.6 | 6.5 | AGPL-3.0 / Enterprise |
| YOLO12n | One-stage, attention-centric YOLO | [41] | Ultralytics 8.4.118 | 640 | 2.6 | 7.6 | AGPL-3.0 / Enterprise |
| YOLO26n | One-stage, end-to-end YOLO | [42] | Ultralytics 8.4.118 | 640 | 2.4 | 5.4 | AGPL-3.0 / Enterprise |
| SSDLite320 MobileNetV3-Large | One-stage SSDLite; MobileNetV3-Large backbone | [43,44] | TorchVision 0.22.1+cu126 | 320 | 3.440 | 0.58 | BSD-3-Clause |
| EfficientDet-D0 | One-stage EfficientDet; EfficientNet-B0 backbone and BiFPN | [45] | effdet 0.4.1 | 512 | 3.88 | 2.54 | Apache-2.0 |
| Faster R-CNN MobileNetV3-Large-320 FPN | Two-stage Faster R-CNN; MobileNetV3-Large backbone and FPN | [44,46] | TorchVision 0.22.1+cu126 | 320 | 19.386 | 0.72 | BSD-3-Clause |

† Parameters and GFLOPs are published specifications for the corresponding reference/pretrained detector variants at the stated reference input size and are not measurements of the 14-class models fine-tuned in this study. For Ultralytics models, these values correspond to the published fused-model specifications.
‡ Software license refers to the implementation used in this study and does not supersede separate terms that may apply to pretrained weights, training datasets, or downstream deployment.

**S1 Video. Prototype RRWIT interactive identification and research-assistance workflow.** The video demonstrates the user-facing RRWIT website, including image upload and live image capture for automated weevil detection and species identification, together with an exploratory conversational artificial intelligence interface for information related to RRWIT and the research program. The conversational component was not systematically evaluated and is not part of the detection benchmark.

## References

[1] Oberprieler RG, Marvaldi AE, Anderson RS. Weevils, weevils, weevils everywhere. Zootaxa. 2007;1668(1):491–520. doi:10.11646/zootaxa.1668.1.24.

[2] McKenna DD, Sequeira AS, Marvaldi AE, Farrell BD. Temporal lags and overlap in the diversification of weevils and flowering plants. Proc Natl Acad Sci U S A. 2009;106(17):7083–7088. doi:10.1073/pnas.0810618106.

[3] Perkin LC, Cohen ZP, Sim SB, Geib SM, Childers AK, Smith TPL, et al. A chromosome level reference genome for the pecan weevil, Curculio caryae. Sci Data. 2026;13:706. doi:10.1038/s41597-026-07030-8.

[4] Harris MK, Jackman JA. Pecan arthropod management. In: Wood BW, Payne JA, editors. Pecan husbandry: challenges and opportunities. First National Pecan Workshop Proceedings. Washington, DC: U.S. Department of Agriculture, Agricultural Research Service; 1991. ARS-96. pp. 6–15.

[5] Harris M, Ree B, Cooper J, Jackman J, Young J, Lacewell R, et al. Economic impact of pecan integrated pest management implementation in Texas. J Econ Entomol. 1998;91(5):1011–1020. doi:10.1093/jee/91.5.1011.

[6] Mulder PG Jr, Harris MK, Grantham RA. Biology and management of the pecan weevil (Coleoptera: Curculionidae). J Integr Pest Manag. 2012;3(1):A1–A9. doi:10.1603/IPM10027.

[7] Tedders WL, Wood BW. A new technique for monitoring pecan weevil emergence (Coleoptera: Curculionidae). J Entomol Sci. 1994;29(1):18–30.

[8] Greenberg SM, Adamczyk JJ Jr, Armstrong JS. Principles and practices of integrated pest management on cotton in the Lower Rio Grande Valley of Texas. In: Larramendy ML, Soloneski S, editors. Integrated pest management and pest control-current and future tactics. Rijeka: InTech; 2012. pp. 3–34. doi:10.5772/31467.

[9] U.S. Department of Agriculture, Animal and Plant Health Inspection Service. Cotton pests [Internet]. Riverdale (MD): USDA APHIS; [cited 2026 Sep 5]. Available from: https://www.aphis.usda.gov/plant-pests-diseases/cotton-pests

[10] Alves AN, Souza WSR, Borges DL. Cotton pests classification in field-based images using deep residual networks. Comput Electron Agric. 2020;174:105488. doi:10.1016/j.compag.2020.105488.

[11] Toscano-Miranda R, Aguilar J, Hoyos W, Caro M, Trebilcok A, Toro M. Different transfer learning approaches for insect pest classification in cotton. Appl Soft Comput. 2024;153:111283. doi:10.1016/j.asoc.2024.111283.

[12] Spiesman BJ, Gratton C, Gratton E, Hines H. Deep learning for identifying bee species from images of wings and pinned specimens. PLoS ONE. 2024;19(5):e0303383. doi:10.1371/journal.pone.0303383.

[13] Liu L, Ouyang W, Wang X, Fieguth P, Chen J, Liu X, et al. Deep learning for generic object detection: a survey. Int J Comput Vis. 2020;128:261–318. doi:10.1007/s11263-019-01247-4.

[14] De Cesaro Junior T, Rieder R. Automatic identification of insects from digital images: a survey. Comput Electron Agric. 2020;178:105784. doi:10.1016/j.compag.2020.105784.

[15] Wang QJ, Zhang SY, Dong SF, Zhang GC, Yang J, Li R, et al. Pest24: a large-scale very small object data set of agricultural pests for multi-target detection. Comput Electron Agric. 2020;175:105585. doi:10.1016/j.compag.2020.105585.

[16] Jiao L, Dong S, Zhang S, Xie C, Wang H. AF-RCNN: an anchor-free convolutional neural network for multi-categories agricultural pest detection. Comput Electron Agric. 2020;174:105522. doi:10.1016/j.compag.2020.105522.

[17] Zhang W, Huang H, Sun Y, Wu X. AgriPest-YOLO: a rapid light-trap agricultural pest detection method based on deep learning. Front Plant Sci. 2022;13:1079384. doi:10.3389/fpls.2022.1079384.

[18] Tang Z, Lu J, Chen Z, Qi F, Zhang L. Improved Pest-YOLO: real-time pest detection based on efficient channel attention mechanism and transformer encoder. Ecol Inform. 2023;78:102340. doi:10.1016/j.ecoinf.2023.102340.

[19] Chen JW, Lin WJ, Cheng HJ, Hung CL, Lin CY, Chen SP. A smartphone-based application for scale pest detection using multiple-object detection methods. Electronics. 2021;10(4):372. doi:10.3390/electronics10040372.

[20] Sittinger M, Uhler J, Pink M, Herz A. Insect detect: an open-source DIY camera trap for automated insect monitoring. PLoS ONE. 2024;19(4):e0295474. doi:10.1371/journal.pone.0295474.

[21] Mostafa V, Sozol MSS, Mondal MRH. LeafDet: a lightweight and interpretable deep learning framework for tomato leaf disease detection. PLoS ONE. 2026;21(5):e0349501. doi:10.1371/journal.pone.0349501.

[22] Jocher G, Qiu J. Ultralytics YOLO11. Version 11.0.0 [software]. 2024. Available from: https://github.com/ultralytics/ultralytics

[23] Quinn M. Weevils of Texas [Internet]. Texas Entomology; [cited 2026 Sep 5]. Available from: https://www.texasento.net/TXWeevils.html

[24] Franz NM, Chamorro ML, Fleming D, Setliff GP, O’Brien CW. Weevils of North America (WoNA) [dataset]. Ag Data Commons; 2017 [cited 2026 Sep 5]. Available from: https://agdatacommons.nal.usda.gov/articles/dataset/Weevils_of_North_America_WoNA_/24852279

[25] iNaturalist. iNaturalist [Internet]. [cited 2026 Sep 5]. Available from: https://www.inaturalist.org/

[26] BugGuide. Coleoptera: Curculionidae [Internet]. Iowa State University; [cited 2026 Sep 5]. Available from: https://bugguide.net/node/view/15740

[27] Rife TW, Poland JA. Field Book: an open-source application for field data collection on Android. Crop Sci. 2014;54(4):1624–1627. doi:10.2135/cropsci2013.08.0579.

[28] Clark JA, contributors. Pillow: Python Imaging Library (Fork) [software]. [cited 2026 Sep 5]. Available from: https://pillow.readthedocs.io/

[29] Liu S, Zeng Z, Ren T, Li F, Zhang H, Yang J, et al. Grounding DINO: marrying DINO with grounded pre-training for open-set object detection. In: Computer Vision-ECCV 2024. Cham: Springer; 2025. pp. 38–55. doi:10.1007/978-3-031-72970-6_3.

[30] Dwyer B, Nelson J, Hansen T, et al. Roboflow. Version 1.0 [software]. 2026. Available from: https://roboflow.com/

[31] Lin TY, Maire M, Belongie S, Hays J, Perona P, Ramanan D, et al. Microsoft COCO: common objects in context. In: Fleet D, Pajdla T, Schiele B, Tuytelaars T, editors. Computer Vision-ECCV 2014. Cham: Springer; 2014. pp. 740–755. doi:10.1007/978-3-319-10602-1_48.

[32] Abid A, Abdalla A, Abid A, Khan D, Alfozan A, Zou J. Gradio: hassle-free sharing and testing of ML models in the wild. arXiv [Preprint]. 2019. arXiv:1906.02569.

[33] Hugging Face. Hugging Face Spaces [Internet]. [cited 2026 Sep 5]. Available from: https://huggingface.co/spaces

[34] Gildenblat J, contributors. PyTorch library for CAM methods [software]. GitHub; 2021. Available from: https://github.com/jacobgil/pytorch-grad-cam

[35] Selvaraju RR, Cogswell M, Das A, Vedantam R, Parikh D, Batra D. Grad-CAM: visual explanations from deep networks via gradient-based localization. In: 2017 IEEE International Conference on Computer Vision. 2017. pp. 618–626. doi:10.1109/ICCV.2017.74.

[36] Muhammad MB, Yeasin M. Eigen-CAM: Class activation map using principal components. In: 2020 International Joint Conference on Neural Networks. 2020. pp. 1–7. doi:10.1109/IJCNN48605.2020.9206626.

[37] Adebayo J, Gilmer J, Muelly M, Goodfellow I, Hardt M, Kim B. Sanity checks for saliency maps. Adv Neural Inf Process Syst. 2018;31.

[38] Woo S, Park J, Lee JY, Kweon IS. CBAM: Convolutional Block Attention Module. In: Proceedings of the European Conference on Computer Vision. 2018. pp. 3–19. doi:10.1007/978-3-030-01234-2_1.

[39] Jocher G. Ultralytics YOLOv5 [software]. 2020. doi:10.5281/zenodo.3908559. Available from: https://github.com/ultralytics/yolov5

[40] Jocher G, Chaurasia A, Qiu J. Ultralytics YOLOv8 [software]. 2023. Available from: https://github.com/ultralytics/ultralytics

[41] Tian Y, Ye Q, Doermann D. YOLOv12: attention-centric real-time object detectors. arXiv [Preprint]. 2025. arXiv:2502.12524. doi:10.48550/arXiv.2502.12524.

[42] Jocher G, Qiu J, Liu M, Lyu S, Akyon FC, Kalfaoglu ME. Ultralytics YOLO26: unified real-time end-to-end vision models. arXiv [Preprint]. 2026. arXiv:2606.03748. doi:10.48550/arXiv.2606.03748.

[43] Liu W, Anguelov D, Erhan D, Szegedy C, Reed S, Fu CY, et al. SSD: Single Shot MultiBox Detector. In: Computer Vision-ECCV 2016. Cham: Springer; 2016. pp. 21–37. doi:10.1007/978-3-319-46448-0_2.

[44] Howard A, Sandler M, Chu G, Chen LC, Chen B, Tan M, et al. Searching for MobileNetV3. In: Proceedings of the IEEE/CVF International Conference on Computer Vision. 2019. pp. 1314–1324. doi:10.1109/ICCV.2019.00140.

[45] Tan M, Pang R, Le QV. EfficientDet: scalable and efficient object detection. In: Proceedings of the IEEE/CVF Conference on Computer Vision and Pattern Recognition. 2020. pp. 10781–10790. doi:10.1109/CVPR42600.2020.01079.

[46] Ren S, He K, Girshick R, Sun J. Faster R-CNN: towards real-time object detection with region proposal networks. IEEE Trans Pattern Anal Mach Intell. 2017;39(6):1137–1149. doi:10.1109/TPAMI.2016.2577031.

[47] Bai J, Lu F, Zhang K, et al. ONNX: Open Neural Network Exchange [software]. GitHub; 2019. Available from: https://github.com/onnx/onnx

[48] ONNX Runtime developers. ONNX Runtime [software]. Available from: https://onnxruntime.ai/

[49] Burke HR. Studies on the genus Anthonomus in North and Central America (Coleoptera: Curculionidae) II. The subgenus Anthonomorphus Dietz. Coleopt Bull. 1964;18(1):7–17. doi:10.5962/p.372349.

[50] Whitehead DR, Chamorro ML, Anderson RS. An illustrated key to the species of Curculio Linnaeus (Coleoptera: Curculionidae) of North America east of the Mississippi River. Proc Entomol Soc Wash. 2018;120(3):616–641. doi:10.4289/0013-8797.120.3.616.

[51] Ali F, Qayyum H, Iqbal MJ. Faster-PestNet: a lightweight deep learning framework for crop pest detection and classification. IEEE Access. 2023;11:104016–104027. doi:10.1109/ACCESS.2023.3317506.

[52] Dong Q, Sun L, Han T, Cai M, Gao C. PestLite: a novel YOLO-based deep learning technique for crop pest detection. Agriculture. 2024;14(2):228. doi:10.3390/agriculture14020228.

[53] Khan A, Malebary SJ, Dang LM, Binzagr F, Song HK, Moon H. AI-enabled crop management framework for pest detection using visual sensor data. Plants. 2024;13(5):653. doi:10.3390/plants13050653.

[54] Xiong P, Zhang C, He L, Zhan X, Han Y. Deep learning-based rice pest detection research. PLoS ONE. 2024;19(11):e0313387. doi:10.1371/journal.pone.0313387.

[55] Liu J, Yu E, Li Y, Zhao Y, Mao B. YOLO-DCPG: a lightweight architecture with dual-channel pooling gated attention for intensive small-target agricultural pest detection. Front Plant Sci. 2025;16:1716703. doi:10.3389/fpls.2025.1716703.

[56] Joshi A, Guevara D, Earles M. Standardizing and centralizing datasets for efficient training of agricultural deep learning models. Plant Phenomics. 2023;5:0084. doi:10.34133/plantphenomics.0084.

[57] Sinanc Terzi D. Effect of different weight initialization strategies on transfer learning for plant disease detection. Plant Pathol. 2024;73:2325–2343. doi:10.1111/ppa.13997.

[58] Vasconcelos C, Birodkar VN, Dumoulin V. Proper reuse of image classification features improves object detection. In: Proceedings of the IEEE/CVF Conference on Computer Vision and Pattern Recognition. 2022. pp. 13628–13637. doi:10.1109/CVPR52688.2022.01326.

[59] Yu C, Xie J, Tony FJA. BGM-YOLO: an accurate and efficient detector for detecting plant disease. PLoS ONE. 2025;20(5):e0322750. doi:10.1371/journal.pone.0322750.

[60] Song Y, An K, Wang L, Zhou B. Multi-scale closed-loop tuning via spatial frequency collaborative sensitivity for rice leaf disease detection. PLoS ONE. 2026;21(6):e0351727. doi:10.1371/journal.pone.0351727.

[61] Karthik RP, Murugesan G, Ballaji HK, Anitha J. UAV-based RGB and multispectral mango leaf disease detection with benchmarking of YOLOv5 to YOLOv10 and SeqOpt-optimised YOLOv8 for real-time edge deployment. PLoS ONE. 2026;21(5):e0349855. doi:10.1371/journal.pone.0349855.

[62] Borianne P, Theveny F, Cabrera-Bosquet L, Lammoglia S-K. CocoaDeep: a preliminary study of the performance sensitivity to datasets of Faster RCNN, YOLO and transformer networks for cocoa pod detection. PLoS ONE. 2026;21(7):e0351657. doi:10.1371/journal.pone.0351657.

[63] Valan M, Makonyi K, Maki A, Vondracek D, Ronquist F. Automated taxonomic identification of insects with expert-level accuracy using effective feature transfer from convolutional networks. Syst Biol. 2019;68(6):876–895. doi:10.1093/sysbio/syz014.

[64] Wu X, Zhan C, Lai YK, Cheng MM, Yang J. IP102: a large-scale benchmark dataset for insect pest recognition. In: Proceedings of the IEEE/CVF Conference on Computer Vision and Pattern Recognition. 2019. doi:10.1109/CVPR.2019.00899.

[65] Schneider S, Taylor GW, Kremer SC, Fryxell JM. Getting the bugs out of AI: advancing ecological research on arthropods through computer vision. Ecol Lett. 2023;26(7):1247–1258. doi:10.1111/ele.14239.

[66] Marais GC, Stratton IC, Johnson AJ, Hulcr J. Progress in developing a bark beetle identification tool. PLoS ONE. 2025;20(6):e0310716. doi:10.1371/journal.pone.0310716.

[67] Chen J, Kao SH, He H, Zhuo W, Wen S, Lee CH, Chan SHG. Run, don’t walk: chasing higher FLOPS for faster neural networks. In: Proceedings of the IEEE/CVF Conference on Computer Vision and Pattern Recognition. 2023. pp. 12021–12031. doi:10.1109/CVPR52729.2023.01157.

[68] Deng X, Qi L, Liu Z, Liang S, Gong K, Qiu G. Weed target detection at seedling stage in paddy fields based on YOLOX. PLoS ONE. 2023;18(12):e0294709. doi:10.1371/journal.pone.0294709.

[69] Lu D, Wang Y. MAR-YOLOv9: a multi-dataset object detection method for agricultural fields based on YOLOv9. PLoS ONE. 2024;19(10):e0307643. doi:10.1371/journal.pone.0307643.

[70] Qin Y, Liu Q, Liu H, Zhang H, Ji Z, Ganchev I. Lightweight real-time detectors of apple-leaf diseases operating on embedded devices. PLoS ONE. 2026;21(7):e0352500. doi:10.1371/journal.pone.0352500.

[71] Jacob B, Kligys S, Chen B, Zhu M, Tang M, Howard A, et al. Quantization and training of neural networks for efficient integer-arithmetic-only inference. In: Proceedings of the IEEE/CVF Conference on Computer Vision and Pattern Recognition. 2018. pp. 2704–2713. doi:10.1109/CVPR.2018.00286.

[72] Wang Z, Ding T, Liang S, Cui H, Gao X. Workpiece surface defect detection based on YOLOv11 and edge computing. PLoS ONE. 2025;20:e0327546. doi:10.1371/journal.pone.0327546.

[73] Chen Y, Sun H, Tian L, Yang Y, Wang S, Wang T. Detecting infrared UAVs on edge devices through lightweight instance segmentation. PLoS ONE. 2025;20(8):e0330074. doi:10.1371/journal.pone.0330074.

[74] Bjerge K, Alison J, Dyrmann M, Frigaard CE, Mann HMR, Hoye TT. Accurate detection and identification of insects from camera trap images with deep learning. PLOS Sustain Transform. 2023;2(3):e0000051. doi:10.1371/journal.pstr.0000051.

[75] He K, Gkioxari G, Dollar P, Girshick R. Mask R-CNN. In: Proceedings of the IEEE International Conference on Computer Vision. 2017. pp. 2961–2969. doi:10.1109/ICCV.2017.322.

[76] Graving JM, Chae D, Naik H, Li L, Koger B, Costelloe BR, et al. DeepPoseKit, a software toolkit for fast and robust animal pose estimation using deep learning. eLife. 2019;8:e47994. doi:10.7554/eLife.47994.

[77] Kirillov A, He K, Girshick R, Rother C, Dollar P. Panoptic segmentation. In: Proceedings of the IEEE/CVF Conference on Computer Vision and Pattern Recognition. 2019. pp. 9404–9413. doi:10.1109/CVPR.2019.00963.

[78] Weyler J, Magistri F, Marks E, Chong YL, Sodano M, Roggiolani G, et al. PhenoBench: a large dataset and benchmarks for semantic image interpretation in the agricultural domain. IEEE Trans Pattern Anal Mach Intell. 2024;46(12):9583–9594. doi:10.1109/TPAMI.2024.3419548.

